# Development of *Corynebacterium glutamicum* as a monoterpene production platform

**DOI:** 10.1101/2023.10.31.565027

**Authors:** Bridget A. Luckie, Meera Kashyap, Allison N. Pearson, Yan Chen, Yuzhong Liu, Luis E. Valencia, Alexander Carrillo Romero, Graham A. Hudson, Xavier B. Tao, Bryan Wu, Christopher J. Petzold, Jay D. Keasling

**Affiliations:** Joint BioEnergy Institute, 5885 Hollis Street, Emeryville, CA 94608, USA; Biological Systems & Engineering Division, Lawrence Berkeley National Laboratory, Berkeley, CA 94720, USA; Department of Molecular and Cell Biology, University of California, Berkeley, CA 94720, USA; Department of Plant and Microbial Biology, University of California, Berkeley, CA 94720, USA; Institute for Quantitative Biosciences, University of California, Berkeley, CA 94720, USA; Joint Program in Bioengineering, University of California, Berkeley/San Francisco, CA 94720, USA; Department of Chemical and Biomolecular Engineering, University of California, Berkeley, CA 94720, USA; The Novo Nordisk Foundation Center for Biosustainability, Technical University of Denmark, Denmark; Center for Synthetic Biochemistry, Institute for Synthetic Biology, Shenzhen Institutes for Advanced Technologies, Shenzhen, China

**Keywords:** Monoterpene biosynthetic production, Geraniol, Citronellol, Eucalyptol, Linalool, Citral, Citronellic acid, *Corynebacterium glutamicum*

## Abstract

Monoterpenes are commonly known for their role in the flavors and fragrances industry and are also gaining attention for other uses like insect repellant and as potential renewable fuels for aviation. *Corynebacterium glutamicum,* a Generally Recognized as Safe microbe, has been a choice organism in industry for the annual million ton-scale bioproduction of amino acids for more than 50 years; however, efforts to produce monoterpenes in *C. glutamicum* have remained relatively limited. In this study, we report a further expansion of the *C. glutamicum* biosynthetic repertoire through the development and optimization of a mevalonate-based monoterpene platform. In the course of our plasmid design iterations, we increased flux through the mevalonate-based bypass pathway, measuring isoprenol production as a proxy for monoterpene precursor abundance and demonstrating the highest reported titers in *C. glutamicum* to date at nearly 1500 mg/L. Our designs also evaluated the effects of backbone, promoter, and GPP synthase homolog origin on monoterpene product titers. Monoterpene production was further improved by disrupting competing pathways for isoprenoid precursor supply and by implementing a biphasic production system to prevent volatilization. With this platform, we achieved 321.1 mg/L of geranoids, 723.6 mg/L of 1,8-cineole, and 227.8 mg/L of linalool. Furthermore, we determined that *C. glutamicum* first oxidizes geraniol through an aldehyde intermediate before it is asymmetrically reduced to citronellol. Additionally, we demonstrate that the aldehyde reductase, AdhC, possesses additional substrate promiscuity for acyclic monoterpene aldehydes.

**Highlights:**

- Design of a mevalonate-based monoterpene production platform in *C. glutamicum*
- Highest production titers of geranoids, eucalyptol, and linalool reported in *C. glutamicum* to date
- Identification of citronellal as an intermediate in the reduction of geraniol to citronellol by *C. glutamicum*

## 1. Introduction

Isoprenoids, also known as terpenes or terpenoids, comprise the largest class of natural products with more than 55,000 identified compounds found across all domains of life. Composed of one or more five-carbon isoprene units, isoprenoids are diverse in both structure and function with significant utility in the flavors, fragrances, materials, insect repellant, biofuels, and therapeutics industries (Keasling et al. 2021; Mosquera et al. 2021; Beller, Lee, and Katz 2015; Soares-Castro, Soares, and Santos 2020; Kozioł et al. 2014). Isoprenoids are biosynthetically derived from the condensation of isopentenyl diphosphate (IPP) and dimethylallyl diphosphate (DMAPP), most commonly produced through either the methylerythritol 4-phosphate (MEP) pathway or the mevalonate-dependent (MVA) pathway. Some of the highest monoterpene titers (grams per liter quantities) to date have been achieved through heterologous, MVA-based production in *Escherichia coli* and augmented MVA-based production in *Saccharomyces cerevisia*e (Jiang et al. 2021; Wang et al. 2021; Zhu et al. 2021; X. Chen, Zhang, and Lindley 2020). While monoterpene production has been expanded to other microbial chassis beyond *E. coli* and *S. cerevisiae* with impressive success (Kirby et al. 2021; Hoshino et al. 2020), efforts to produce monoterpenes in *Corynebacterium glutamicum* thus far remain relatively limited in both the titers and breadth of monoterpene products (~15 mg/L or less) (Li, Xu, and Lu 2021; Kang et al. 2014).

*C. glutamicum*, an industrial workhorse responsible for the million ton-scale production of amino acids each year, has increasingly been leveraged to produce natural products such as polyketides, organic acids, polyphenols, and alcohols (Cankar, Henke, and Wendisch 2023; Zha et al. 2023; Wolf et al. 2021). The intrinsic characteristics of *C. glutamicum* have enabled it to become a pillar of biotechnology success with particular importance to the consumer products and food additive industry. More specifically, *C. glutamicum* is a fast-growing, endotoxin-free, generally regarded as safe (GRAS) bacterium capable of growing to high cell densities while retaining metabolic activity and withstanding variations typical of industrial growth conditions. Moreover, it possesses superior recombinant protein expression due to its minimal protease activity as well as the ability to assimilate multiple carbon sources in a monoauxic growth state (Ray et al. 2022). However, despite its keys advantages and impressive natural product repertoire, only a limited set of isoprenoids have been produced in *C. glutamicum (Moser and Pichler 2019; Heider and Wendisch 2015; Park and Woo 2022; Henke et al. 2018; Frohwitter et al. 2014; H. Lim, Park, and Woo 2020; Li, Xu, and Lu 2021)*. Each of these efforts relied upon the overexpression of key components [typically deoxyxylulose 5-phosphate synthase (*dxs*) and isopentenyl diphosphate isomerase (*idi*)] within the native MEP pathway, which is subject to endogenous regulation that may limit the isoprenyl precursors, IPP and DMAPP (Banerjee and Sharkey 2014; Li, Xu, and Lu 2021; Kang et al. 2014; Di et al. 2023). To overcome these potential limitations, (Sasaki et al. 2019) leveraged a heterologous MVA-based, episomal bypass pathway to produce isoprenol at titers over 1000 mg/L, providing a promising foundation toward high production for other isoprenoids.

Herein we report the development of an episomal, mevalonate-based platform to expand the natural product repertoire of *C. glutamicum* to achieve the highest titers of geraniol-derived monoterpenes (geranoids), linalool, and 1,8-cineole (eucalyptol) in this organism to date. Furthermore, we investigated the mechanism by which geraniol is asymmetrically reduced to citronellol in *C. glutamicum* and determined that geraniol first proceeds through an aldehyde intermediate before being reduced; however, the enzymatic mechanism of reduction remains unknown.

**Figure 1.**
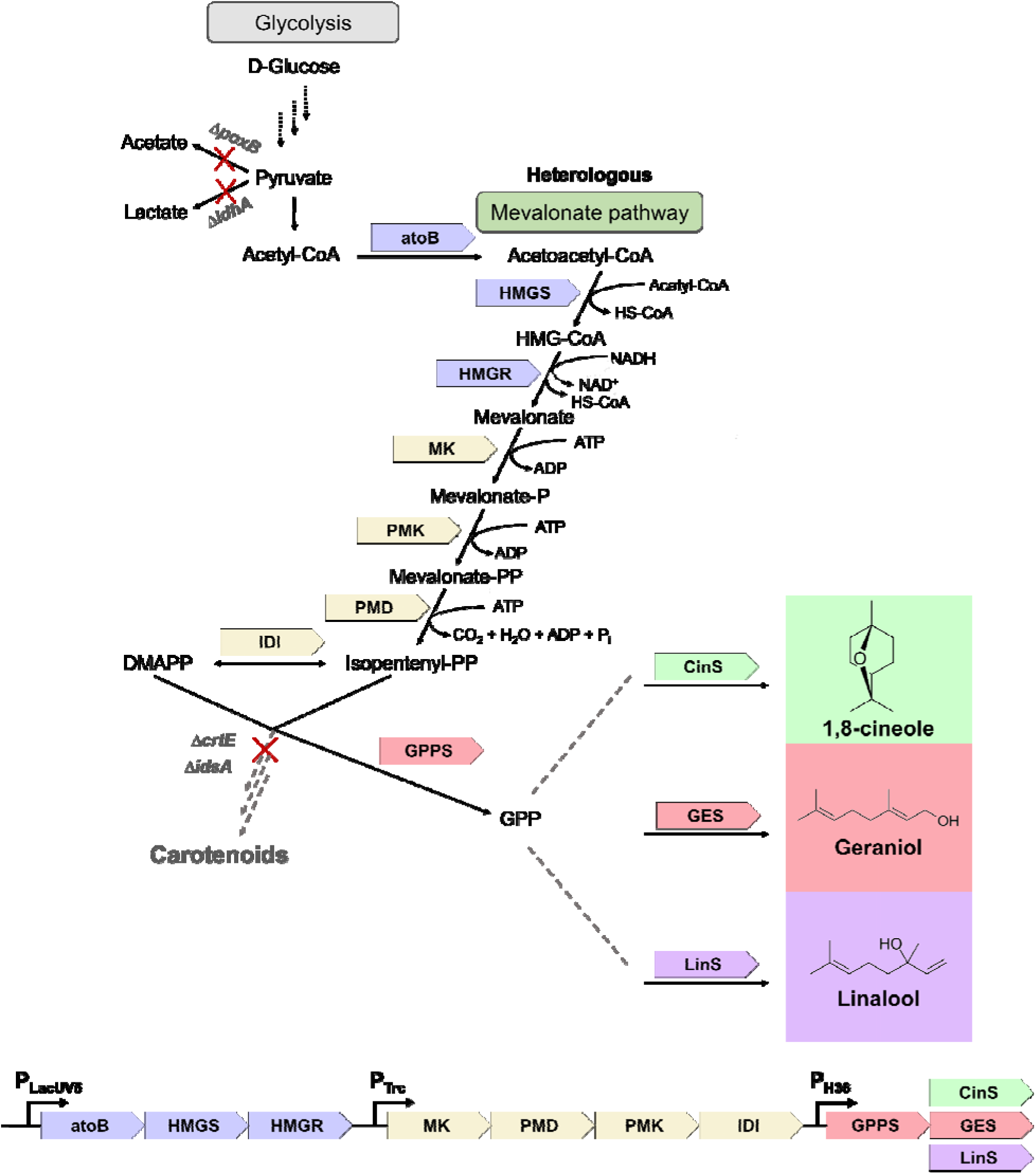
Pathway and engineering strategy. The heterologous mevalonate pathway for monoterpene production is depicted. Targeted pathways are indicated with a red “X” and the corresponding gene deletion. Abbreviations: *poxB*, pyruvate dehydrogenase; *ldhA*, L-lactate dehydrogenase; *crtE,* geranylgeranyl pyrophosphate synthase; *idsA*, geranylgeranyl pyrophosphate synthase; *atoB*, acetyl-CoA acetyltransferase; *HMGS*, hydroxymethylglutaryl-CoA synthase; *HMGR*, 3-hydroxy-3-methylglutaryl-CoA reductase; *MK*, mevalonate kinase; PMK, phosphomevalonate kinase; *PMD*, phosphomevalonate decarboxylase; HMG-CoA, 3-hydroxy-3-methyl-glutaryl-coenzyme A; *IDI*, isopentenyl diphosphate isomerase; *GPPS*, geranyl diphosphate synthase; *CinS*, cineole synthase; *GES*, geraniol synthase; *LinS*, linalool synthase.

## 2. Materials and Methods

### 2.1. Chemicals and reagents

All chemicals and reagents of molecular biology or analytical grade were purchased from Sigma-Aldrich (St. Louis, MO) unless indicated otherwise. Deep well plates used for routine cell cultivation were from Axygen (Union City, CA) and were sealed with a gas-permeable AeraSeal adhesive film (Excel Scientific, Victorville, CA). Brain heart infusion broth (BHI) was purchased from Neogen (Lansing, MI) and CGXII media (without biotin or protocatechuic acid) was purchased from Teknova (Hollister, CA). Miller Luria broth (LB) was purchased from Merck KGaA (Darmstadt, DE).

### 2.2. Strain and plasmid construction and verification

All strains and plasmid sequences developed or used in this study can be found in Table 1 and requested at https://public-registry.jbei.org/folders/808. The names and additional information for genes targeted in this study can be found in Table S1 and sequences can be accessed at https://img.jgi.doe.gov/. Integrated DNA Technologies, Inc (San Diego, CA) synthesized all oligonucleotides primers and double-stranded DNA fragments. Primers utilized for amplifying genomic DNA to sequence and confirm gene deletions can be found in Table S2. All polymerase chain reactions (PCR) used for routine cloning were conducted using Q5 High Fidelity DNA polymerase 2X Master Mix (New England BioLabs, Ipswich, MA). Plasmids were constructed using isothermal DNA assembly with 40-60 bp overhangs (NEBuilder HiFi DNA Assembly Master Mix, New England BioLabs, Ipswich, MA) and transformed into high-efficiency NEB 10-beta competent *E. coli* (New England BioLabs, Ipswich, MA). Phusion® High-Fidelity PCR Master Mix with HF Buffer (New England BioLabs) was used to screen *C. glutamicum* for gene deletions and amplicons of the target size were sent to Genewiz/Azenta (South San Francisco, CA) for validation. To acquire genomic DNA from *C. glutamicum* for PCR, stationary phase cultures were boiled in 80% dimethyl sulfoxide (DMSO) at 99 °C for 10 minutes to lyse cells. Plasmid sequences were verified by Primoridium Labs (Monrovia, CA). Codon optimizations for stated genes were performed by the Gensmart™ Codon Optimization Tool (GenScript, Piscataway, NJ).

**Table 1.** Strains and plasmids.

| Strain | Registry # | Description | Selection | Reference |
| --- | --- | --- | --- | --- |
| JBEI-7936 | JPUB_014552 | *C. glutamicum* JBEI 1.1.2, biotin auxotroph | Nx<sup>R</sup> |  |
| PL | JPUB_013248 | JBEI-7936 Δ*poxB* Δ*ldhA* | Suc<sup>R</sup> Kan<sup>S</sup> | (Sasaki et al. 2019) |
| PLC | JPUB_022481 | JBEI-7936 Δ*poxB* Δ*ldhA* Δ*crtE* | Suc<sup>R</sup> Kan<sup>S</sup> | This study |
| PLI | JPUB_022484 | JBEI-7936 Δ*poxB* Δ*ldhA* Δ*idsA* | Suc<sup>R</sup> Kan<sup>S</sup> | This study |
| PLCI | JPUB_022487 | JBEI-7936 Δ*poxB* Δ*ldhA* Δ*crtE* Δ*idsA* | Suc<sup>R</sup> Kan<sup>S</sup> | This study |
| IPK | JPUB_022506 | Δ*poxB* Δ*ldhA* containing pJBEI-19625 | Suc<sup>R</sup> Kan<sup>S</sup> | (Sasaki et al. 2019) |
| IPWT | JPUB_022507 | JBEI-7936 containing pECXT202 | Suc<sup>R</sup> Tet<sup>R</sup> | This study |
| IPT | JPUB_022476 | Δ*poxB* Δ*ldhA* containing pECXT202 | Suc<sup>R</sup> Tet<sup>R</sup> | This study |
| WT-44 | JPUB_022509 | JBEI-7936 containing pBAL44 | Suc<sup>R</sup> Tet<sup>R</sup> | This study |
| PL-44 | JPUB_022474 | ΔpoxB ΔldhA containing pBAL44 | Suc<sup>R</sup> Kan<sup>R</sup> | This study |
| PL-114 | JPUB_022472 | ΔpoxB ΔldhA containing pBAL114 | Suc<sup>R</sup> Tet<sup>R</sup> | This study |
| PLC-114 | JPUB_022482 | ΔpoxB ΔldhA ΔcrtE + pBAL114 | Suc<sup>R</sup> Tet<sup>R</sup> | This study |
| PLI-114 | JPUB_022494 | ΔpoxB ΔldhA ΔidsA + pBAL114 | Suc<sup>R</sup> Tet<sup>R</sup> | This study |
| PLCI-114 | JPUB_022490 | ΔpoxB ΔldhA ΔcrtE ΔidsA + pBAL114 | Suc<sup>R</sup> Tet<sup>R</sup> | This study |
| PL-118 | JPUB_022478 | ΔpoxB ΔldhA containing pBAL118 | Suc<sup>R</sup> Kan<sup>R</sup> | This study |
| PL-121 | JPUB_022477 | ΔpoxB ΔldhA containing pBAL121 | Suc<sup>R</sup> Kan<sup>R</sup> | This study |
| PL-218 | JPUB_022495 | ΔpoxB ΔldhA containing pBAL218 | Suc<sup>R</sup> Tet<sup>R</sup> | This study |
| PLC-218 | JPUB_022497 | ΔpoxB ΔldhA ΔcrtE + pBAL218 | Suc<sup>R</sup> Tet<sup>R</sup> | This study |
| PLI-218 | JPUB_022485 | ΔpoxB ΔldhA ΔidsA + pBAL218 | Suc<sup>R</sup> Tet<sup>R</sup> | This study |
| PLCI-218 | JPUB_022488 | ΔpoxB ΔldhA ΔcrtE ΔidsA + pBAL218 | Suc<sup>R</sup> Tet<sup>R</sup> | This study |
| PL-230 | JPUB_022504 | ΔpoxB ΔldhA containing pBAL230 | Suc<sup>R</sup> Tet<sup>R</sup> | This study |
| PL-231 | JPUB_022505 | ΔpoxB ΔldhA containing pBAL231 | Suc<sup>R</sup> Tet<sup>R</sup> | This study |
| PLCI-230 | JPUB_022510 | ΔpoxB ΔldhA ΔcrtE ΔidsA containing pBAL230 | Suc<sup>R</sup> Tet<sup>R</sup> | This study |
| PLCI-231 | JPUB_022511 | ΔpoxB ΔldhA ΔcrtE ΔidsA containing pBAL231 | Suc<sup>R</sup> Tet<sup>R</sup> | This study |
| BL-1 | JPUB_022514 | Δ3088 | Suc<sup>R</sup> | This study |
| BL-2 | JPUB_022513 | Δ3092 | Suc<sup>R</sup> | This study |
| BL-3 | JPUB_022512 | Δ3088 Δ3092 | Suc<sup>R</sup> | This study |
| BL-4 | JPUB_022532 | Δ225 Δ1733 Δ2308 Δ2399 Δ2854 Δ3088 Δ3092 | Suc<sup>R</sup> | This study |
| BL-5 | JPUB_022498 | ΔadhC | Suc<sup>R</sup> | This study |
| BL-6 | JPUB_022515 | ΔadhA | Suc<sup>R</sup> | This study |
| BL-7 | JPUB_022516 | Δ225 Δ1733 Δ2308 Δ2399 Δ2854 Δ3088 Δ3092 ΔadhC | Suc<sup>R</sup> | This study |
| BL-8 | JPUB_022517 | $\Delta 2844$ | Suc<sup>R</sup> | This study |
| BL-9 | JPUB_022518 | $\Delta 2844 \Delta adhC$ | Suc<sup>R</sup> | This study |
| BL-10 | JPUB_023382 | JBEI-7936 containing pECXT-pSyn | Tet<sup>R</sup> | This study |
| BL-11 | JPUB_023384 | $\Delta adhC$ containing pECXT-pSyn | Tet<sup>R</sup> | This study |
| BL-12 | JPUB_023383 | JBEI-7936 containing pBAL272 | Tet<sup>R</sup> | This study |
| BL-13 | JPUB_023385 | $\Delta adhC$ containing pBAL272 | Tet<sup>R</sup> | This study |
| **Plasmid** |  |  |  |  |
|  | pJBEI-2600 | pEC-XK99E, *E. coli* - *C. glutamicum* shuttle expression vector containing pGA1, oriV, Ptrc | Kan<sup>R</sup> | (Kirchner and Tauch 2003) |
|  | pJBEI-19625 | pTE202 pEC-XK99E-AK-IP-bypass-S. pomeroyi HMGR (substitution), *E. coli*-*C. glutamicum* shuttle expression vector containing pGA1 origin, oriV, p<sub>LacUV5</sub> p<sub>Trc</sub>lacI<sup>q</sup> | Kan<sup>R</sup> | Sasaki et al., 2019 |
|  |  | pECXT-pSyn, *E. coli*-*C. glutamicum* shuttle expression vector containing pGA1 origin, Tet<sup>R</sup>, oriV | Tet<sup>R</sup> | Henke et al., 2021 |
|  | JPUB_022519 | pECXT202, pECXT-IP-bypass-S. Pomeroyi HMGR (pECXT-pSyn/pTE202 derivative) | Tet<sup>R</sup> | This study |
|  | JPUB_022520 | pBAL44 pTE202-PMK-IDI-H36-ERG20<sup>ww</sup>-ObGEScP | Kan<sup>R</sup> | This study |
|  | JPUB_022521 | pBAL114 pECXT202-PMK-IDI-H36-ERG20<sup>ww</sup>-ObGEScP | Tet<sup>R</sup> | This study |
|  | JPUB_022522 | pBAL118 pTE202-PMK-IDI-H36-AgGPPS-ObGEScP | Kan<sup>R</sup> | This study |
|  | JPUB_022524 | pBAL121 pTE202-PMK-IDI-pSyn-AgGPPS-ObGEScP | Kan<sup>R</sup> | This study |
|  | JPUB_022526 | pBAL218 pECXT202-PMK-IDI-H36-AgGPPS-ObGEScP | Tet<sup>R</sup> | This study |
|  | JPUB_022527 | pBAL230 pECXT202-PMK-IDI-H36-AgGPPS-CinSco | Tet<sup>R</sup> | This study |
|  | JPUB_022528 | pBAL231 pECXT202-PMK-IDI-H36-AgGPPS-LinSco | Tet<sup>R</sup> | This study |
|  | JPUB_022530 | pBAL16 pk18-Δ225 | Suc<sup>S</sup> Kan<sup>R</sup> | This study |
|  | JPUB_022534 | pBAL23 pk18-Δ1733 | Suc<sup>S</sup> Kan<sup>R</sup> | This study |
|  | JPUB_022537 | pBAL24 pk18-Δ2308 | Suc<sup>S</sup> Kan<sup>R</sup> | This study |
|  | JPUB_022540 | pBAL25 pk18-Δ2399 | Suc<sup>S</sup> Kan<sup>R</sup> | This study |
|  | JPUB_022542 | pBAL26 pk18-Δ2854 | Suc<sup>S</sup> Kan<sup>R</sup> | This study |
|  | JPUB_022545 | pBAL27 pk18-Δ3088 | Suc<sup>S</sup> Kan<sup>R</sup> | This study |
|  | JPUB_022548 | pBAL28 pk18-Δ3092 | Suc<sup>S</sup> Kan<sup>R</sup> | This study |
|  | JPUB_022551 | pBAL98 pk18mobsacB-ΔcrtE | Suc<sup>S</sup> Kan<sup>R</sup> | This study |
|  | JPUB_022553 | pBAL99 pk18mobsacB-ΔidsA | Suc<sup>S</sup> Kan<sup>R</sup> | This study |
|  | JPUB_022555 | pBAL220 pk18mobsacB-ΔadhC | Suc<sup>S</sup> Kan<sup>R</sup> | This study |
|  | JPUB_022557 | pBAL221 pk18mobsacB-ΔadhA | Suc<sup>S</sup> Kan<sup>R</sup> | This study |
|  | JPUB_022559 | pBAL241 pk18mobsacB-Δ2844 | Suc<sup>S</sup> Kan<sup>R</sup> | This study |
|  | JPUB_023386 | pBAL272 pECXT-P<sub>LacUV5</sub>-*adhC* | Tet<sup>R</sup> | This study |

### 2.3. Media, buffer, and solution compositions

Brain Heart Infusion (BHI): 17.5 g/L Brain Heart infusion solids (Porcine), 10 g/L Tryptose, 2 g/L glucose, 5 g/L NaCl, 2.5 g/L Na_2_HPO_4_. CGXII minimal media (Sasaki et al. 2019): 40 g/L D-glucose, 20 g/L (NH4)2SO4, 5 g/L urea, 1 g/L KH2PO4, 1 g/L K2HPO4, 0.25 g/L MgSO4·7H2O, 10 mg/L CaCl2, 10 mg/L FeSO4·7H2O, 10 mg/L MnSO4·H2O, 1 mg/L ZnSO4·7H2O, 0.2 mg/L CuSO4·5H2O, 0.02 mg/L NiCl2·6H2O, 0.2 mg/L biotin, 30 mg/L 3,4-dihydroxybenzoic acid, and 21 g/L 3-morpholinopropanesulfonic acid (MOPS); pH 7.0. BHI media for electrocompetent *C. glutamicum* cell preparation (BHISGIT) (H. C. Lim et al. 2019): 37 g/L BHI powder, 91g/L sorbitol, 25 g/L glycine, 0.4 g/L isonicotinic acid hydrazide, 1 mL/L Tween 80. Lysis buffer: 50 mM NaCl, 25 mM HEPES (pH 7-7.5), 5% (v/v) glycerol. Trace metals solution (100X): 1 g/L FeSO_4_.7H_2_O, 1 g/L MnSO_4_.H_2_O, 100 mg/L ZnSO_4_.7H_2_O, 20 mg/L CuSO_4_.5H_2_O, 2 mg/L NiCl_2_.6H_2_O.

### 2.4. Culture conditions

Routine culturing of *E. coli* was performed in LB at 37°C. Routine culturing of *C. glutamicum* was performed in BHI at 30°C. When necessary, *E. coli* and *C. glutamicum* were respectively cultivated in medium containing 50 or 25 μg/mL of kanamycin sulfate or 5 μg/mL of tetracycline hydrochloride. Production of monoterpenes was conducted in CGXII minimal media (Sasaki et al. 2019). Routine adaptation into CGXII involved diluting a stationary phase culture in BHI 1:10 into CGXII twice over 48 hours.

### 2.5. Preparation and electroporation of electrocompetent *C. glutamicum* cells

*C. glutamicum* was made electrocompetent through a previously described method (Lim et al. 2019). Briefly, *C. glutamicum* was grown overnight in a starter culture that was subsequently diluted 1:100 into BHISGIT media, grown 14-18 hours to an OD_600_ of 0.5 - 0.8, and then chilled on ice for 1 hour. Cells were harvested by centrifugation at 3000 × g at 4°C for 10 minutes. Cells were then washed 3 times with 50 mL ice cold 10% glycerol, resuspended in ice cold 10% glycerol to a final OD_600_ of 20, and stored as 50 microliter aliquots at −80°C until use. Cells were electroporated using a 1 mm cuvette, 200 ng of vector, and a GenePulser Xcell Electroporator (Bio-Rad, Hercules, CA) at 200 Ω, 25 μF, and 19.5 kV/cm. Competent cells were subsequently transferred into 1 mL of BHI, heat-shocked for 6 min at 46 °C, and allowed to recover for 2.5 hours at 30 °C before plating on BHI agar plates containing the appropriate antibiotic.

### 2.6. Monoterpene toxicity time course assay

Following routine adaptation to CGXII, stationary cells were back diluted to an OD_600_ of 0.1 in 0.5 mL of CGXII followed by the exogenous addition of the related monoterpene in a Falcon 48-well tissue culture-treated cell culture plate (Corning, NY) sealed with Breathe Easy gas-permeable sealing membrane (Diversified Biotech, Dedham, MA). Optical density measurements were conducted at 600 nm every 10 minutes over 96 hours in a SpectraMax M2e plate reader (Molecular Devices, San Jose, CA) with orbital shaking at 30 °C.

### 2.7. Isoprenol and monoterpene production and extraction

All strains were initially inoculated into a starter culture of BHI media containing the appropriate antibiotic in a 24 deep-well plate and incubated at 30 overnight with shaking at 200 rpm. The cultures were subsequently passaged twice (1:10) every 24 hours in CGXII media containing the appropriate antibiotic and then inoculated into 5 mL of CGXII at an OD_600_ of 0.1. For experiments where an organic overlay was utilized, squalane was added at 10 percent of the culture media volume 12-24 hours after induction. The strains utilized to generate the data depicted in and preceding Figure 3b were induced at an OD_600_ of ~0.8 with 0.5 mM isopropyl β-D-1-thiogalactopyranoside (IPTG) in test tube cultures containing 5 mL of CGXII. All subsequent production experiments were conducted under the same conditions with the exception of the induction time where cells were induced at an OD_600_ of 0.1 with the same concentration of IPTG due to increased product titers and improved experimental ease (Fig. S3).

### 2.8. Monoterpene biotransformation and volatilization experiments

Starter cultures of relevant *C. glutamicum* strains were grown overnight in BHI to stationary phase. For monoterpene feeding experiments carried out in BHI, each strain was back diluted to a starting OD_600_ of 2 and grown for 24 hours at 30 on a rotary shaker at 200 rpm in the presence of 100 mg/L of the respective monoterpene. For monoterpene feeding experiments carried out in CGXII, cells were subjected to two rounds of adaptation in CGXII. The adapted cells were then back diluted to an OD_600_ of 2 and grown on a rotary shaker at 200 rpm in the presence of 100 mg/L of the respective monoterpene. For measuring volatilization, 100 mg/L of the respective monoterpene was added to sterile 5 mL CGXII or BHI. All monoterpenes were diluted in DMSO or ethanol. For the biotransformation experiment submitted for proteomics analysis, cells were inoculated in fresh BHI at an OD_600_ of 2 and either 5 µL of 100% ethanol or 100 mg/L citral (2.25% solution in ethanol, *v/v*) was applied. Cultures were incubated as described above. For the experiments where AdhC was overexpressed, cells were adapted in CGXII as previously described and induced with 0.5 mM IPTG at an OD_600_ of 0.1 in 5mL cultures. DMSO containing 100 mg/L of citral was added 16 hours following induction and cultures were harvested 24 hours after the introduction of citral.

### 2.9. *In vitro* assay of crude lysate to determine cofactor requirement(s)

From an overnight culture, 75 mL of stationary phase cells were added to 450 mL BHI and incubated for 24 hours. The cells were subsequently harvested at 3000 × *g*, washed with the lysis buffer, and spun down at 3000 × g for 20 min. The cell pellet was stored at −80 until lysis. For lysis, the cell pellet was resuspended 100 mL of lysis buffer and 2 mL aliquot was collected and stored on ice. The remaining entire volume was subjected to six rounds of pressure homogenization at 17,000 psi at 4. A 2 mL aliquot of the crude cell lysate was collected and stored on ice. The cell lysate was subsequently centrifuged at 40,000 × *g* at 4, and the supernatant was decanted to collect the pellet and soluble protein fractions. The fractions were subsequently used or snap frozen and stored at −80 until use. For the assay, the soluble protein fraction (or whole or crude lysed cells where indicated) was diluted 1 to 5 to a final volume of 200 uL in lysis buffer containing either 5 mM NAD^+^, NADH, NADP^+^, NADPH, FMN, and/or 1X trace metals solution. The reaction was initiated with the addition of 100 mg/L geraniol or citral (where indicated) diluted in DMSO and incubated at 30 for 2 hours. The reaction was quenched with an equal volume of ethyl acetate and processed for GC-MS analysis as discussed below.

### 2.10. Isoprenol and monoterpene analysis by GC-MS/GC-FID

Cultures were harvested as previously described (Sasaki et al. 2019). Briefly, 200-300 μL culture underwent liquid:liquid extraction with an equal volume of ethyl acetate containing either 25 mg/L nerolidol or, in the case of linalool-containing samples, 30 mg/L 1-butanol as an internal standard for 15 minutes at 3000 rpm in a Vortex Genie 2 (Scientific Industries, INC, NY, USA). After centrifugation at 14,500 × *g*, ~150 uL of the organic phase was transferred to an Agilent glass insert in a glass vial. For samples utilizing squalane as an overlay, 1 mL of culture was harvested and pelleted at 14,500 × *g* for 2.5 minutes and 10 uL of the organic phase was added to 990 uL of ethyl acetate in a glass vial. Next, 1 µL was injected into either an Agilent Intuvo 9000 gas chromatograph mass spectrometer (GC-MS) or Agilent 8890 gas chromatograph flame ionization detector (GC-FID). The GC-MS was equipped with an Agilent DB-WAX UI column either 15 m × 0.25 mm, 0.25 um in length and the GC-FID was equipped with a DB-WAX UI column 15 m × 0.32 mm × 0.25 um in length. Following injection into the GC-MS at a flow rate of 1 mL/min, the oven was held at 60 for 1 min, followed by a ramp at 15/min to 150, then 10/min to 200, and 30/min to 240 where it was held for 1 minute. For detection of geranic and citronellic acid by GC-MS, the oven was instead held at 60 for 1 minute, followed by a ramp at 15/min to 150, then 10/min to 250 where it was held for 0.5 minute. Following injection into the GC-FID at a flow rate of 2.2 mL/min, the oven was held at 40 for 1 minute, followed by a ramp of 15/min to 100, and a ramp of 30/min until 230 where it was held for 1.5 minutes. In all cases, the inlet temperature was held at 250. Analytical grade standards were used to generate standard curves for quantification of analytes and peak confirmation. The product concentrations reported from samples containing squalane as an organic overlay were calculated according to the aqueous culture volume.

### 2.11. Determination of sugar in the culture medium

Glucose was measured by high-performance liquid chromatography (HPLC) using an Aminex HPX-87H column (BioRad Laboratories, USA) on an Agilent Technologies 1200 series HPLC with a refractive index detector. At each timepoint, 500 uL of culture was harvested by centrifugation and the supernatant was quenched with an equal volume of ice-cold methanol. Samples were stored at −80 until use. The mobile phase (4 mM sulfuric acid) was run at a flow rate of 0.6 mL/min and the column temperature was set to 60. The injection volume was 10 μL in all cases. The peak areas and respective concentrations were interpolated from a standard calibration curve created using authentic standards.

### 2.12. Synthesis of deuterated (D_2_)-geraniol

The synthetic procedure was adapted from (Duhamel et al. 2016). A solution of geranic acid (0.159 g, 0.95 mmol) in diethyl ether (1 mL) was added dropwise to a slurry of lithium aluminum deuteroxide (0.08 g, 1.9 mmol) in diethyl ether (5 mL) at 0 °C. The resulting mixture was warmed to room temperature and left to stir for 2 h. Wet diethyl ether (5 mL) was added slowly and stirred for 10 mins. Saturated aqueous ammonium chloride (5 mL) was added, and extracted with diethyl ether (3 × 10 mL). The combined organic extracts were dried (Na2SO4), and the solvent was removed *in vacuo*. The synthesized D_2_-geraniol was subsequently analyzed by ^1^H NMR (Fig. S7) with the corresponding chemical shifts:

^1^H NMR (400 MHz, CDCl3) δ: 5.42 (1H, t), 5.10 (1H, t), 2.10 (4H, m), 1.68 (6H, s), 1.60 (3H, s).

### 2.13. Proteomics analysis

At the indicated time points, 1 mL of culture was harvested by centrifugation, discarding the supernatant, flash frozen, and stored at −80 until proteomics analysis. Protein was extracted and tryptic peptides were prepared by following established proteomic sample preparation protocol (Y. Chen et al. 2023). Briefly, cell pellets were resuspended in Qiagen P2 Lysis Buffer (Qiagen, Hilden, Germany, Cat.#19052) to promote cell lysis. Proteins were precipitated with addition of 1 mM NaCl and 4 volumetric equivalents of acetone, followed by two additional washes with 80% acetone in water. The recovered protein pellet was homogenized by pipetting mixing with 100 mM ammonium bicarbonate in 20% methanol. Protein concentration was determined by the DC protein assay (BioRad, Hercules, CA). Protein reduction was accomplished using 5 mM tris 2-(carboxyethyl)phosphine (TCEP) for 30 min at room temperature, and alkylation was performed with 10 mM iodoacetamide (IAM; final concentration) for 30 min at room temperature in the dark. Overnight digestion with trypsin was accomplished with a 1:50 trypsin:total protein ratio. The resulting peptide samples were analyzed on an Agilent 1290 UHPLC system coupled to a Thermo Scientific Orbitrap Exploris 480 mass spectrometer for discovery proteomics (Y. Chen, Gin, and Petzold 2022). Briefly, peptide samples were loaded onto an Ascentis® ES-C18 Column (Sigma–Aldrich, St. Louis, MO) and separated with a 10 minute gradient from 98% solvent A (0.1 % FA in H2O) and 2% solvent B (0.1% FA in ACN) to 65% solvent A and 35% solvent B. Eluting peptides were introduced to the mass spectrometer operating in positive-ion mode and were measured in data-independent acquisition (DIA) mode with a duty cycle of 3 survey scans from *m/z* 380 to *m/z* 985 and 45 MS2 scans with precursor isolation width of 13.5 *m/z* to cover the mass range. DIA raw data files were analyzed by an integrated software suite DIA-NN (Demichev et al. 2020). The database used in the DIA-NN search (library-free mode) is *C. glutamicum* JBEI 1.1.2 proteome FASTA sequence from the Joint Genome Institute plus the protein sequences of the heterologous proteins and common proteomic contaminants. DIA-NN determines mass tolerances automatically based on first pass analysis of the samples with automated determination of optimal mass accuracies. The retention time extraction window was determined individually for all MS runs analyzed via the automated optimization procedure implemented in DIA-NN. Protein inference was enabled, and the quantification strategy was set to Robust LC = High Accuracy. Output main DIA-NN reports were filtered with a global FDR = 0.01 on both the precursor level and protein group level. The Top3 method, which is the average MS signal response of the three most intense tryptic peptides of each identified protein, was used to plot the quantity of the targeted proteins in the samples ((Silva et al. 2006; Ahrné et al. 2013).

For our analysis of the proteomics data, we calculated the log_2_-fold difference between the Top3 percent abundance of each protein in the citral condition versus the ethanol control. To generate a heat map of the proteins with a significant fold-difference, we filtered the data so that we excluded all proteins that did not meet the following criteria for at least one of the three timepoints: a standard deviation less than 25% of the Top3 percent abundance value, a log_2_ fold-difference greater than |1|, and a value for the Top3 counts greater than 10^5^. Analysis and plotting was performed using Python 3.9, Seaborn 0.12.2, and Matplotlib 3.7.2.

### Data Availability

The generated mass spectrometry proteomics data have been deposited to the ProteomeXchange Consortium via the PRIDE (Perez-Riverol et al. 2022) partner repository with the dataset identifier PXD044193.

## 3. Results and Discussion

### 3.1. *C. glutamicum* plasmid backbone and expression optimization increases isoprenol and monoterpene production

Several groups have demonstrated that *C. glutamicum* is a viable host for heterologous production of monoterpenes (Kang et al. 2014; Li, Xu, and Lu 2021). These efforts relied on augmentation of the endogenous MEP pathway by overexpressing key enzymes; however, titers were limited. Recent studies have demonstrated a conserved, allosteric regulatory mechanism where 1-deoxy-D-xylulose-5-phosphate synthase (DXS), the rate-limiting MEP enzyme, forms inactive aggregates when IPP and DMAPP levels are high (Banerjee and Sharkey 2014; Di et al. 2023). The subsequent downregulation of the MEP pathway may be a source of the low titers achieved in the previous *C. glutamicum* monoterpene production studies. This limitation was likely circumvented when Sasaki et al. (2019) leveraged a mevalonate-based bypass pathway to produce 1250 mg/L isoprenol (IP) in a *C. glutamicum* strain lacking pyruvate dehydrogenase (encoded by *poxB*) and lactate dehydrogenase (encoded by *ldhA*). Based on this work, we hypothesized that a mevalonate-based platform could be harnessed to improve monoterpene production by decoupling IPP and DMAPP pools from the native MEP regulatory mechanisms. To establish this platform in *C. glutamicum*, we leveraged the IP bypass pathway plasmid (pTE202) with the addition of four additional genes: phosphomevalonate kinase (*ERG8/PMK* from *S. cerevisiae*), isopentenyl diphosphate isomerase (*idi* from *E. coli*), a GPP synthase [*ERG20*^WW^ from *S. cerevisiae,* (Ignea et al. 2014) or truncated *AgGPPS2* from *Abies grandis*, (Burke and Croteau 2002)], and a truncated version of the prototypical monoterpene synthase, geraniol synthase [*ObGEScP* from *Ocimum basilicum* (<u>Denby et al 2018</u>)].

We first constructed a variant of the isoprenol bypass pathway plasmid, swapping kanamycin resistance (pTE202) for tetracycline resistance (pECXT202), and evaluated the isoprenol production from each of these plasmids in the wild-type (WT) and Δ*poxB* Δ*ldhA* strains (Fig. 2a). Surprisingly, the product titers of the) Δ*poxB* Δ*ldhA* strain containing pECXT202 (IPT) increased by 1.28-fold compared to the Δ*poxB* Δ*ldhA* strain containing pTE202 (IPK) (Fig. 2b, 1429.3 mg/L ± 102.9 vs. 1116.5 mg/L ± 85.4, respectively) which was not due to differences in growth (Fig. S2b).

**Figure 2.**
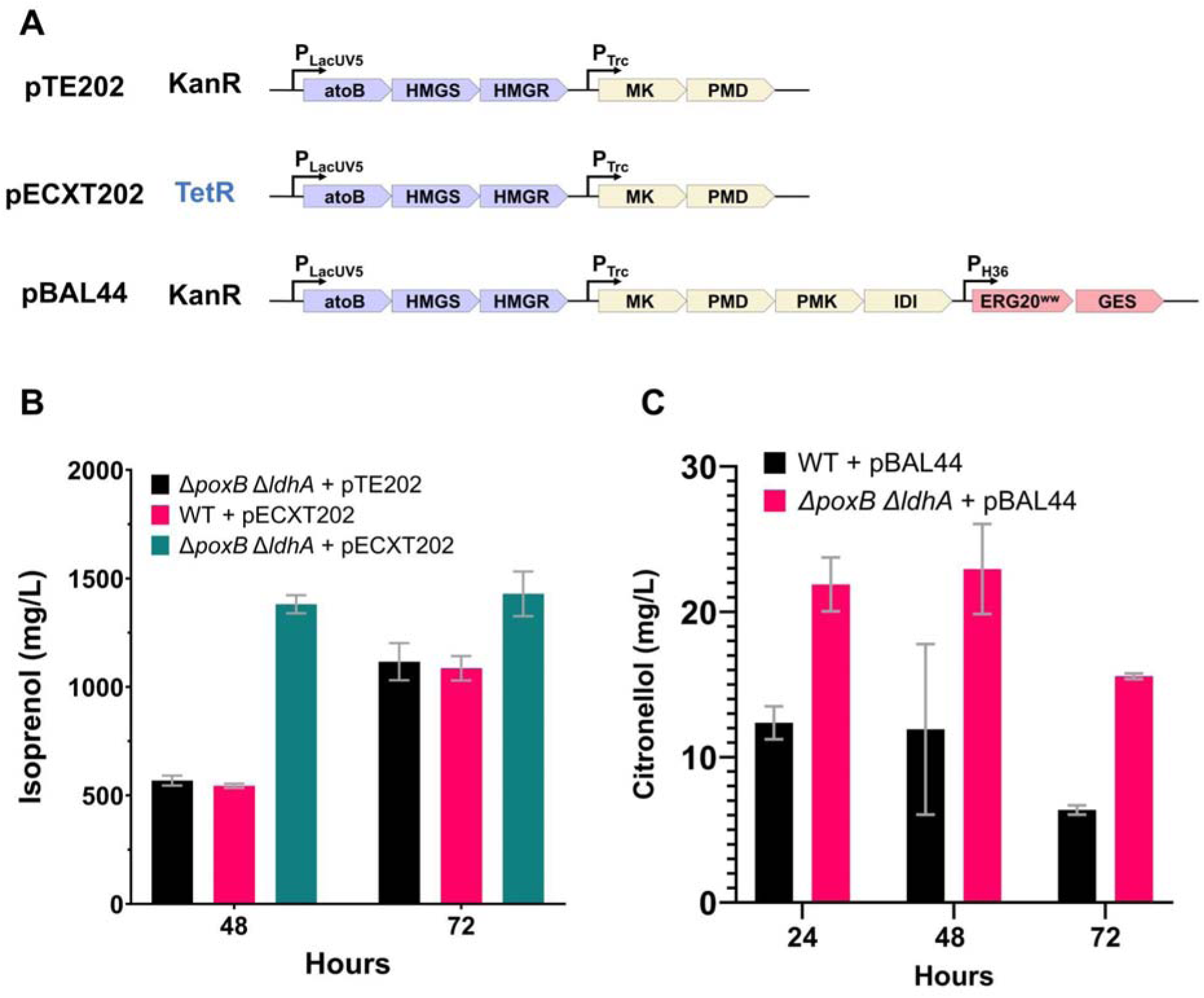
Assessment of strain background and antibiotic resistance on isoprenol and citronellol titers. (a) Schematics of the episomal expression operons for isoprenol and geraniol production. (b) Isoprenol production utilizing the IPK, IPWT, and IPT strains. (c) Citronellol production from the WT-44 and PL-44 strains. All strains were induced with 0.5 mM IPTG at an OD_600_ of 0.8 and harvested at the indicated times following induction.

Because the Δ*poxB* Δ*ldhA* background led to the highest isoprenol titers by reducing acetyl-CoA flux into competing, upstream pathways, we assessed whether this strain background would similarly be suitable for monoterpene production. We tested this strain background by utilizing the KanR plasmid pBAL44, which contains the MVA pathway, the GPP synthase *ERG20^WW^,* and the geraniol synthase *ObGEScP* (Fig. 2a). Interestingly, citronellol was the primary product in the culture medium of this strain (PL-44) with smaller amounts of geraniol and nerol (Fig. 2c). Notably, *C. glutamicum* has not previously been reported to catalyze the asymmetric reduction of geraniol to citronellol, the mechanism of which will be discussed later in this work. The Δ*poxB* Δ*ldhA* strain (PL-44) outperformed WT (WT-44) by almost 2-fold (22.95 mg/L ± 3.09 vs. 11.91 mg/L ± 5.87, Fig. 2c) and thus was selected as the strain background for subsequent experiments.

Next, the TetR analogue of pBAL44 was generated (pBAL114, Fig. 3a), which led to a 1.85-fold improvement in geranoid production compared to pBAL44 and a corresponding increase in pathway expression (Fig. 3a-c). The relative expression of the five proteins shared between the isoprenol (Fig. S1) and monoterpene (Fig. 3b) strains were comparable, indicating the precursor flux was likely not the source limiting monoterpene production. This led us to investigate the relatively low expression of the third operon containing the GPP synthase and geraniol synthase as the bottleneck.

**Figure 3.**
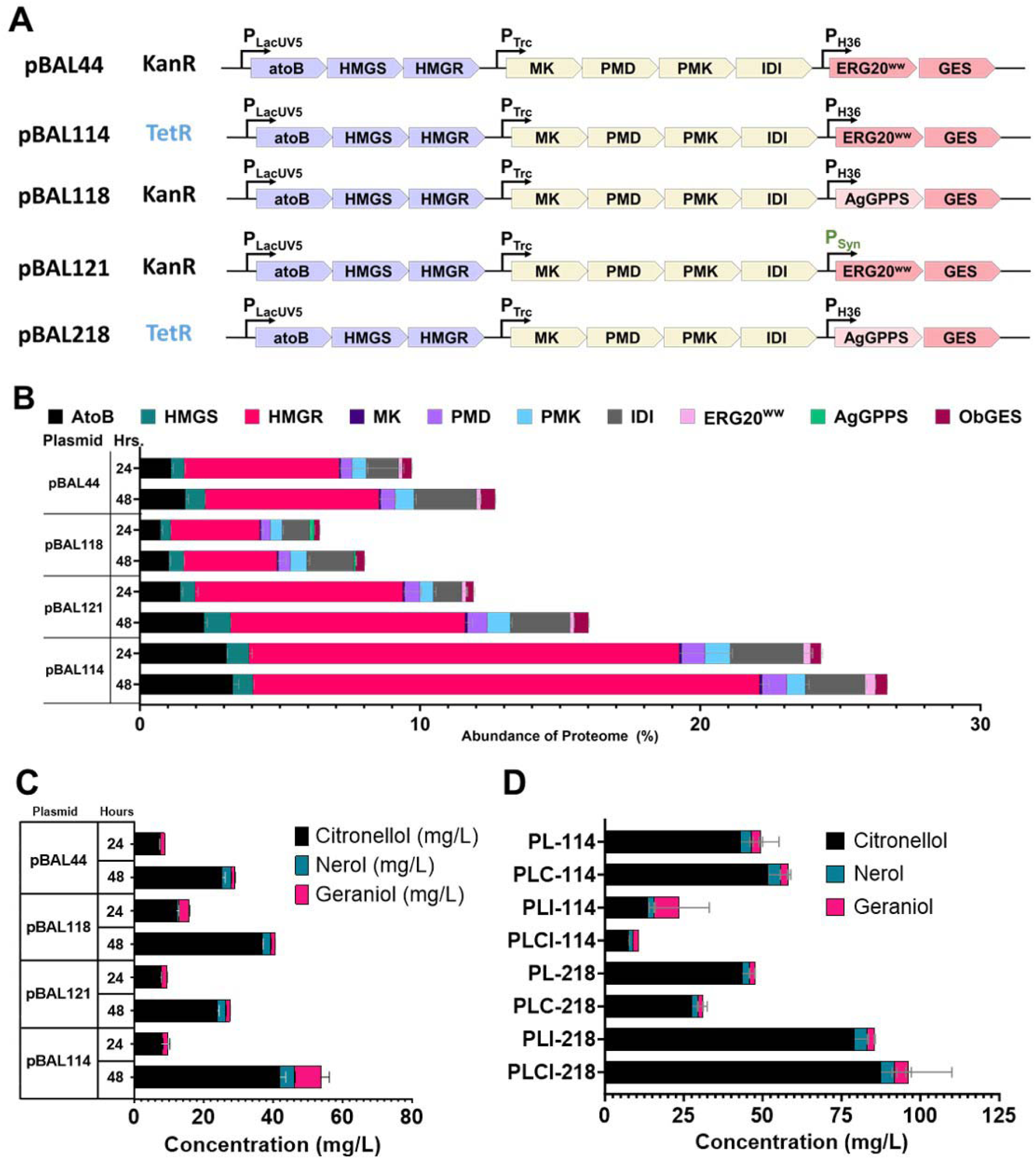
Optimization of operon expression, GPP synthase origin, and metabolite flux. (a) Schematics of the optimization modifications for geraniol production. (b) Expression of pathway proteins which were quantified by using the Top3 “best flyer” quantification method (Silva et al. 2006). (c-d) Geranoid production following expression optimization and strain engineering. (b-c) Plasmids were evaluated in the Δ*poxB* Δ*ldhA* background and were induced at an OD_600_ of 0.8. (d) Culture was induced at an OD_600_ of 0.1. Production was measured at the indicated times after induction (c) and at 60 hours (d).

To tackle this, we first replaced *ERG20^WW^* with the GPP synthase from *A. grandis* (*AgGPPS*) to produce the KanR plasmid, pBAL118, which resulted in a 1.39-fold increase in product titers (40.56 mg/L, ± 0.27) compared to the KanR-pBAL44 (29.08 mg/L, ± 1.02, Fig. 2c) despite similar levels of expression (Fig. 3a-c). Additionally, P_Syn_, a strong synthetic, constitutive promoter (Rytter et al. 2014; Henke, Krahn, and Wendisch 2021), was also chosen in an attempt to improve expression of the third operon containing *ERG20^WW^*and *ObGEScP* to generate the KanR plasmid, pBAL121 (Fig. 3a,c). While protein expression from genes in the first and second operons did slightly increase at 48 hours in strains harboring pBAL121 compared to the original strain (pBAL44), neither the expression of *ERG20^ww^* and *ObGEScP* nor the product titers appreciably increased compared to those produced by the same strain harboring pBAL44 (27.56 mg/L, ± 0.67 mg/L vs. 29.08 mg/L, ± 1.02, Fig. 3a-c). To test if *AgGPPS* in a TetR backbone would synergistically improve titers, we generated PL-218 which surprisingly did not result in an appreciable increase in titers compared to PL-114 (Fig. 3a-c).

### 3.2. Redirection of flux increases monoterpene titers in concert with catalytically efficient GPP synthase

In parallel, we aimed to further increase geranoid titers by reducing downstream consumption of IPP and DMAPP by competing metabolic pathways. Previous efforts to decrease the flux of IPP and DMAPP into competing carotenoid biosynthetic pathways by targeting two geranylgeranyl diphosphate synthases, CrtE and IdsA, led to an increase in valencene and squalene production in *C. glutamicum (Frohwitter et al. 2014; Park and Woo 2022)*. Interestingly, iteratively targeting *crtE* and *idsA* in strains harboring either pBAL114 or pBAL218 resulted in opposite effects on titers (Fig. 3d). While the Δ*poxB* Δ*ldhA* Δ*crtE* strain harboring pBAL114 (PLC-114) yielded a slight increase in titers, Δ*poxB* Δ*ldhA* Δ*crtE* harboring pBAL218 (PLC-218) resulted in a slight decrease (Fig. 3d). Conversely, Δ*poxB* Δ*ldhA* Δ*idsA* and Δ*poxB* Δ*ldhA* Δ*crtE* Δ*idsA* strains harboring pBAL114 (PLI-114 and PLCI-114) led to a decrease in titers where the same strains harboring pBAL218 (PLI-218 and PLCI-218) yielded a 1.77-fold and 2-fold increase over the Δ*poxB* Δ*ldhA* strain, respectively (Fig. 3d). These differences do not appear to be due to variation in biomass or growth rate (Fig. S4). Instead, we suspected that these differences may be due to differences in specific activity or improper folding under production conditions in *C. glutamicum*. ERG20^WW^ may generate GPP less efficiently than AgGPPS, which could lead to a toxic accumulation of IPP and DMAPP when IdsA, the predominant GGPP synthase in C. glutamicum, is not present. Notably, a significantly decreased transformation efficiency was observed when generating PLCI-114, a phenomenon we typically associate with toxicity. Furthermore, these results are in agreement with a previous study where *ERG20^WW^* was leveraged to produce geraniol in *C. glutamicum,* in which targeting *idsA* but not *crtE* led to a decrease in geraniol production (Li, Xu, and Lu 2021). While this prior study achieved a modest 15 mg/L of geraniol, the cooperative deletion of both *crtE* and *idsA* with the use of AgGPPS in PLCI-218 yielded 98.22 mg/L (± 24.44) of geranoid products.

### 3.3 Expanding the monoterpene biosynthetic repertoire of *Corynebacterium glutamicum* to linalool and 1,8-cineole

Once PLCI-218, a strain which efficiently produced geranoids at approximately 100 mg/L was established, we sought to leverage the mevalonate-based platform to produce other monoterpenes by substituting the type of monoterpene synthase. We selected linalool and 1,8-cineole (eucalyptol) as potential products because we found that *C. glutamicum* is tolerant to both at concentrations of 2 g/L or greater (Fig. S2b,c). Furthermore, both have been successfully produced in other microorganisms by harnessing actinobacterial-derived monoterpene synthases (Hoshino et al. 2020; Ferraz et al. 2021; Mendez-Perez et al. 2017). Therefore, we selected and codon-optimized the 1,8-cineole synthase (CinSco) and linalool synthase (LinSco) from *Streptomyces clavuligeris* to construct plasmids pBAL230 and pBAL231 (Fig. 4a), respectively.

**Figure 4.**
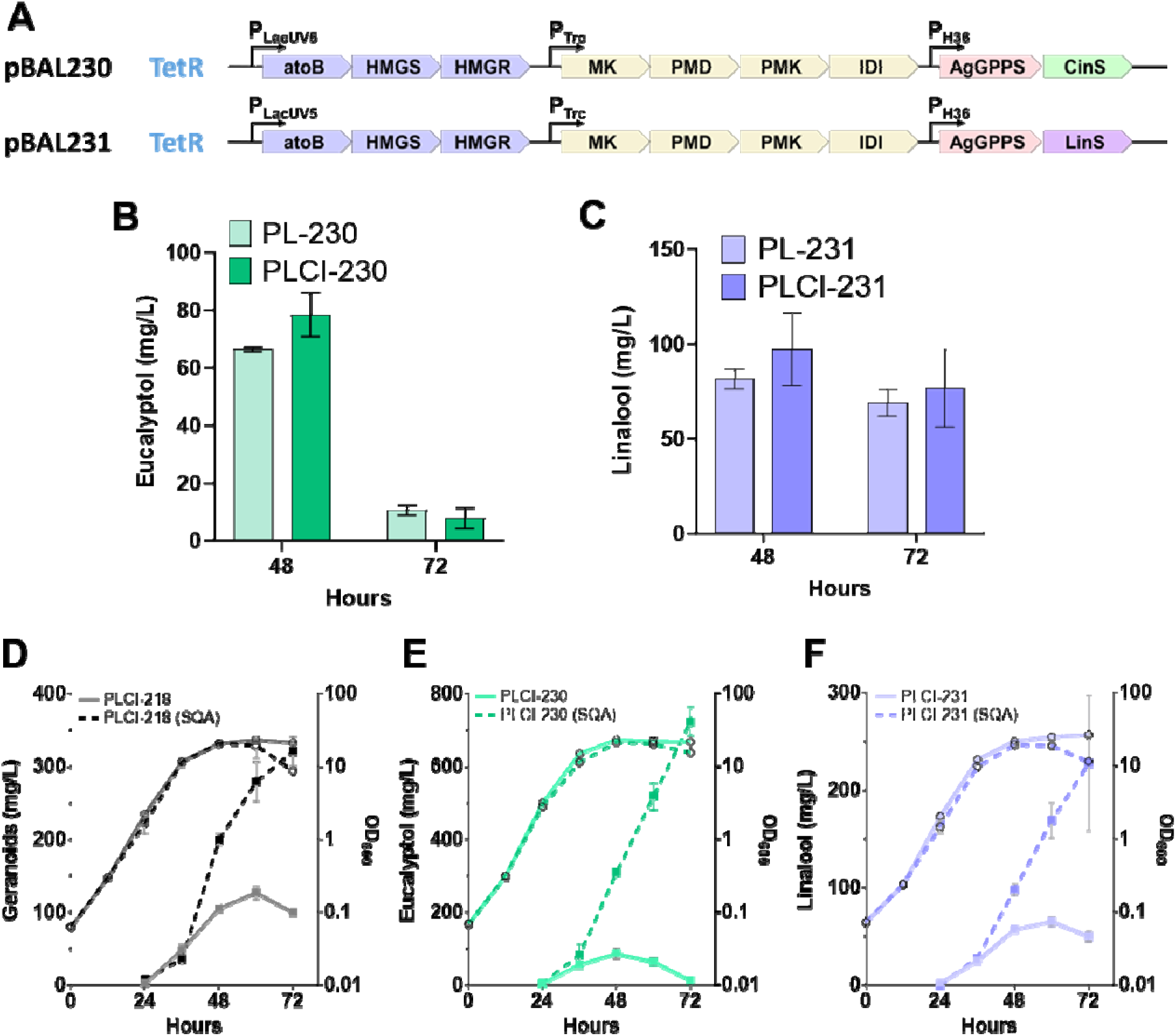
Production and capture of geranoids, eucalyptol, and linalool using a squalane overlay. (a) Schematics of the expression system for eucalyptol (pBAL230) and linalool (pBAL231) production. Production of eucalyptol (b) and linalool (c) from the PL and PLCI strains. Production of monoterpenes (closed squares, d-f) with and without 10% squalane overlay (SQA), added at 12 hours after induction and the corresponding biomass (open circles). All production strains were induced at an OD_600_ of 0.1 and samples were harvested at the indicated times following induction.

Following the success of targeting *crtE* and *idsA* to increase flux in geranoid producing strains, we introduced pBAL230 and pBAL231 into the Δ*poxB* Δ*ldhA* and Δ*poxB* Δ*ldhA* Δ*crtE* Δ*idsA* strains. The Δ*poxB* Δ*ldhA* Δ*crtE* Δ*idsA* strains (PLCI-230 and PLCI-231) outperformed the Δ*poxB* Δ*ldhA* strains (PL-230 and PL-231) harboring both plasmids (Fig. 4b,c); however, the titers for both products decreased after 48 hours, with the effect most pronounced for eucalyptol. This loss indicated that the products were either being catabolized or dissipating into the headspace. To determine the cause, we incubated each of our monoterpene products in sterile media or in the presence of *C. glutamicum.* Eucalyptol was exceedingly volatile and virtually undetectable after 24 hours of incubation in both cell culture and sterile media; however, the loss appeared to be entirely due to volatilization as the rate of eucalyptol loss in media containing *C. glutamicum* was virtually identical to the sterile media control (Fig. S5b). Similarly, linalool was highly volatile in both the cell culture and sterile media, with approximately 80% of the initial amount lost after 24 hours and more than 90% lost after 48 hours (Fig. S5c). Interestingly, citronellol was also volatile and not consumed by *C. glutamicum*, losing approximately 60% of the initial amount after 48 hours (Fig. S5a).

### 3.4. Squalane is a compatible organic layer for trapping geranoids, linalool, and eucalyptol in a biphasic culture system

We sought to identify a strategy to reduce the loss of these volatile monoterpenes. Organic overlays like dodecane, nonane, isopropyl myristate (IPM), heptamethylnonane (HMN), and oleyl alcohol (OA) have been traditionally utilized as part of a two-phase production system for capturing volatile products (Muñoz et al. 2008; Chacón et al. 2019; Liu et al. 2016; George et al. 2015; Ferraz et al. 2021). However, the growth of our strain was inhibited in the presence of dodecane and nonane and product titers declined precipitously (2-fold or more) in the presence of IPM, HMN, and OA (data not shown). Therefore, we tested dodecanol, squalene, and squalane. While dodecanol inhibited growth and squalene proved to be unsuitable for downstream GC-MS analysis due to its column incompatibility, squalane permitted growth and did not reduce titers. With the use of a 10% squalane overlay (v/v) and the Δ*poxB* Δ*ldhA* Δ*crtE* Δ*idsA* strain background, we found that strains PLCI-218, PLCI-230, and PLCI-231 produced 321.1 mg/L (± 20.3 mg/L) geranoids, 723.6 mg/L eucalyptol (± 39.5 mg/L), and 227.8 mg/L linalool (± 69.4 mg/L) (Fig. 4d-f), respectively. PLCI-218 reached a maximal production rate of 13.63 mg L^-1^ h^-1^ between 36 and 48 hours at which point the production rate slowed over the remainder of the experiment (Fig. 4d). Comparatively, PLCI-230 and PLCI-231 maintained their maximal production rate over the course of stationary phase (36 - 72 hours), reaching 17.79 mg L^-1^ h^-1^ and 5.56 mg L-1 h-1, respectively (Fig. 4e-f). It is worth noting that, as the production experiment progresses, cells increasingly co-localize into the squalane layer which is likely the reason for the observed decline in cell density in the biphasic cultures at later time points (Fig. 4d-f), particularly in the geranoid and linalool production strains. This may be indicative of product toxicity caused by an inadequate overlay for monoterpene alcohols, especially given the sequestration of geranoid aldehydes by the overlay which are not apparent in monophasic cultures (Fig. S6). Thus, further examination of organic overlays for use during production of monoterpene alcohols in this host is needed.

### 3.5. *C. glutamicum* asymmetrically reduces geraniol to citronellol through an aldehyde intermediate

Interestingly, the squalane overlay captured significant amounts of geranial and neral (the mixture of which is commercially known as citral), leading us to suspect these aldehydes as intermediates in the reduction of geraniol to citronellol (Fig. S6). However, the generation of multiple products is not ideal for industrial production. As the biotransformation of geraniol to citronellol has not yet been described in any *C. glutamicum* strain, we sought to investigate its mechanism. To first confirm the route of reduction through an aldehyde intermediate, we incubated *C. glutamicum* in the presence of [1,1-D_2_]-geraniol (D_2_-geraniol). The two most probable routes of conversion are (i) direct reduction of geraniol to citronellol leading to D_2_-citronellol or (ii) the formation of an aldehyde intermediate resulting in the loss of one deuterium atom to form D_1_-citronellol (Fig. 5a). Confirming our hypothesis that geraniol was first converted into an aldehyde before being subsequently reduced to citronellol, the major product following incubation with D_2_-geraniol was D_1_-citronellol (Fig. 5b and Fig. S8).

**Figure 5.**
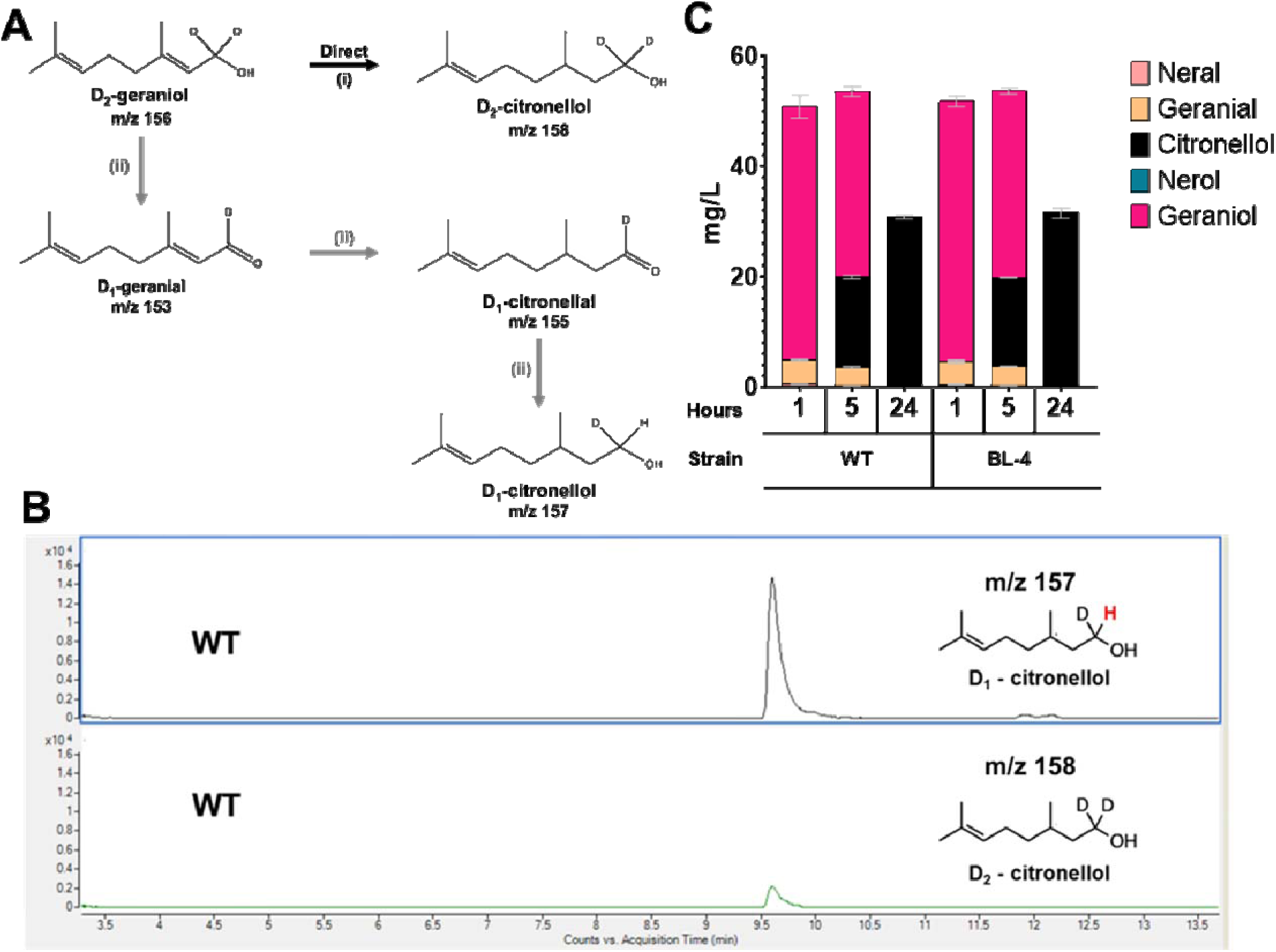
Identifying the mechanism of geraniol reduction to citronellol. (a) Proposed conversion routes of geraniol reduction to citronellol are depicted using deuterium-labeled geraniol. (b) GC-MS chromatogram of isolated parent ions of deuterated citronellol [m/z 157 (D_1-_citronellol) and m/z 158 (D_2_-citronellol)] after 24 hours of incubation in CGXII with *C. glutamicum.* (c) Geranoid products from the biotransformation of 100 mg/L geraniol by WT or BL-4 harvested at the indicated time points.

### 3.6. *C. glutamicum* old yellow enzymes are not responsible for the reduction of geraniol/citral to citronellol/citronellal

Next, we sought to identify the endogenous enzyme(s) responsible for catalyzing the reduction of geraniol/citral. Many alkene reductases belonging to the Old Yellow enzyme family have been shown to catalyze the reduction of geraniol or citral to citronellol in other microbes (Yuan et al. 2011; Ribeaucourt et al. 2022; Richter, Gröger, and Hummel 2011). In *C. glutamicum* JBEI 1.1.2, there are two old yellow enzyme (OYE) homologs (CgI_3088 and CgI_3092) predicted as 2,4-dienoyl-CoA reductase-like NADH-dependent reductases which we selected to target first. Surprisingly, the capacity of strain BL-3 (Δ*3088* Δ*3092*) to reduce geraniol to citronellol was unaffected (Fig. S9). Therefore, we widened our targets to other putative alkene reductases based on their predicted function including four NADPH-dependent 2,4-dienoyl-CoA reductase/sulfur reductase-like enzyme (CgI_225, CgI_1733, CgI_2308, and CgI_2854) and a geranylgeranyl reductase family protein (CgI_2399). Iterative deletion of all seven targeted reductases resulted in a strain (strain BL-4) with no appreciable defect in geraniol reductase activity (Fig. 5c). Additionally, we examined the cofactor requirements for the biotransformation of geraniol to citronellol *in vitro* by using crude cell lysate and found that citronellol only appeared in lysates containing NADPH and improved with the addition of trace metals (Fig. S10). It is therefore unsurprising that the NADH-dependent OYE homologs were not responsible for this reaction.

### 3.7. AdhC acts as an aldehyde reductase on geranoids and its loss leads to accumulation of citronellal and citronellic acid

Next, we aimed to identify the alcohol dehydrogenase/aldehyde reductase(s) acting on the monoterpene alcohols. We selected two targets, AdhC and AdhA, based on their homology to plant geraniol dehydrogenases and the *E. coli* YjgB, YahK, and AdhP/YddN alcohol dehydrogenases which were previously characterized for geraniol dehydrogenase activity (Zhou et al. 2014). In prior studies on *C. glutamicum*, AdhC (also known as FudC) was determined to reduce furfural, and its deletion was leveraged to accumulate aromatic aldehydes and prevent their reduction to their respective alcohols, indicating some substrate promiscuity (Tsuge et al. 2016; Kim et al. 2022). We expected that targeting an enzyme responsible for acting on at least one of the monoterpene alcohols would slow or disrupt the conversion of geraniol to citral and/or citronellal to citronellol. The deletion of *adhC* (strain BL-5) but not *adhA* (strain BL-6) revealed an impact on geraniol reduction to citronellol at early time points (5 hours or earlier) following exogenous addition of geraniol or citral, leading to the slight accumulation of citronellal and decrease in nerol (Fig. 6a,b). While this may suggest that AdhC functions as an aldehyde reductase with a substrate preference for citronellal and neral over geranial, the loss of AdhC ultimately did not prevent the conversion of geraniol to citronellol after 24 hours (Fig. S12b). Interestingly, however, the total concentration of citronellol in the Δ*adhC* strain (BL-5) was more than 75% lower than the wildtype strain after 24 hours incubation with geraniol (Fig. S12b). This suggests that citral or accumulated citronellal may be further oxidized into citronellic acid. Supporting this hypothesis, the Δ*adhC* strain harboring an empty vector (BL-11) accumulated citronellic acid following incubation with citral after 24 hours (Fig. 6e). Furthermore, episomally overexpressing AdhC in either the WT or Δ*adhC* strains incubated with citral resulted in the exclusive accumulation of the geranoid alcohols in nearly identical ratios (Fig. 6d,f). In these strains, nerol and citronellol comprise approximately 40% each of the biotransformed citral whereas geraniol forms approximately 23%. These results collectively demonstrate that AdhC possesses additional substrate promiscuity for acyclic aldehydes and its loss permits accumulated citronellal to become available to an aldehyde dehydrogenase for partial conversion to citronellic acid (Fig. 6g).

**Figure 6.**
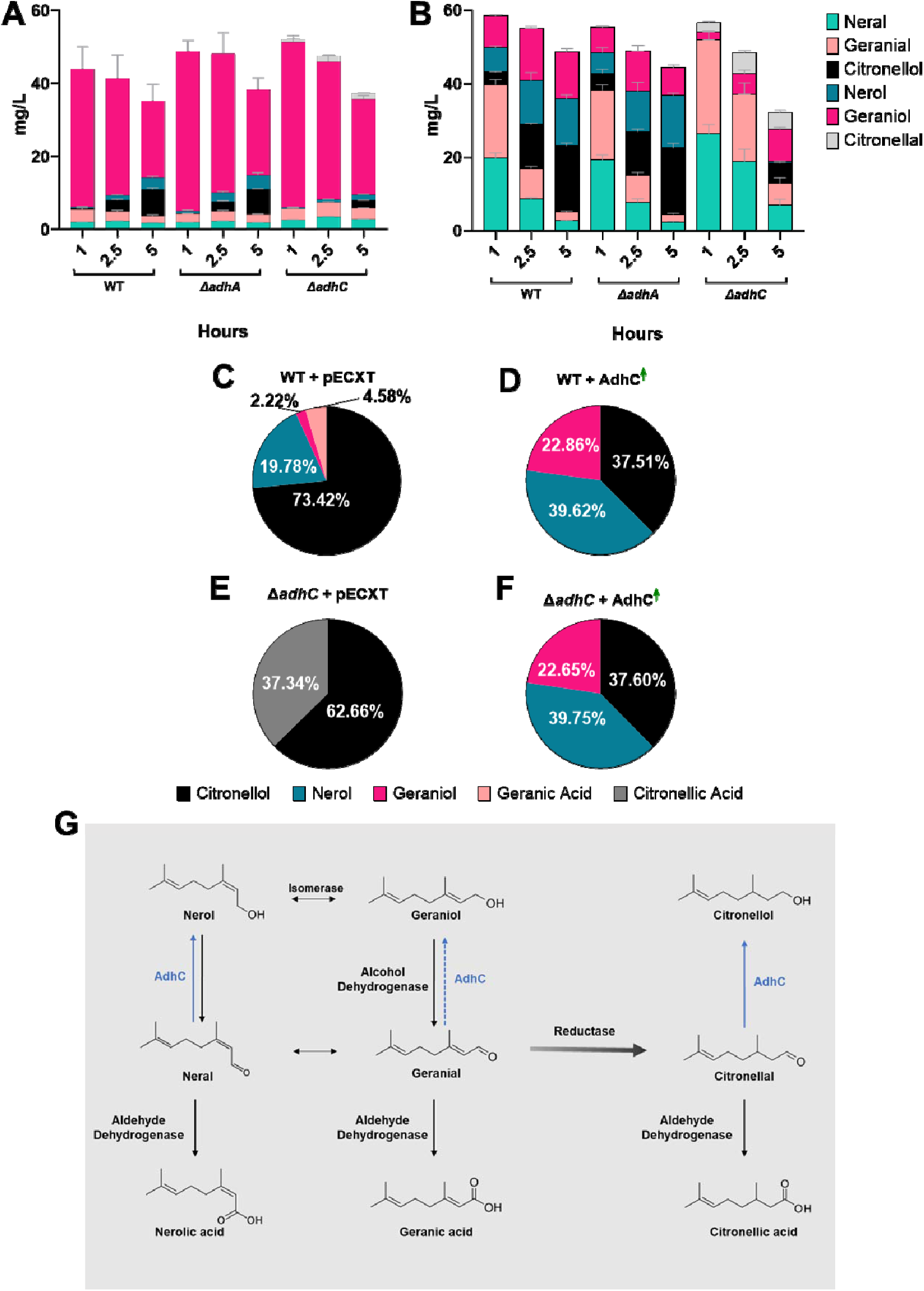
Assessing geraniol oxidation. *C. glutamicum* strains were incubated with 100 mg/L geraniol (a) or citral (b-f) to determine the identity and degree of oxidation and reduction products. The green superscript arrow (d and f) indicates overexpression of AdhC and pECXT (c and e) is an empty vector. The schematic of the proposed route of conversion in *C. glutamicum* and role of AdhC is depicted in (g).

### 3.8. Investigating citral-responsive proteins to determine secondary enzymes induced following the loss of AdhC

In an attempt to identify the aldehyde dehydrogenase or secondary aldehyde reductase oxidizing or reducing citral or citronellal, we investigated differentially expressed proteins following the exogenous addition of citral or the diluent, ethanol (Fig. S11). We defined citral-responsive proteins as proteins that exhibited a more than two-fold difference in percent abundance compared to the ethanol diluent control in at least one time point and had a standard deviation less than 25% of the average fold difference. We then identified citral-responsive proteins that were either unique to or shared between WT and Δ*adhC* (BL-5). The total number of citral-responsive proteins was approximately seven times greater in Δ*adhC* (BL-5) compared to WT (273 vs. 38), and only 17 citral-responsive proteins were shared between the two strains (Fig. S11a). Of these, we identified a putative acyl-coA reductase-like NAD-dependent aldehyde dehydrogenase (CgI_2844) that appeared to be citral-responsive in both strains (Fig. S11b). Because WT and Δ*adhC* containing empty vectors were respectively capable of oxidizing citral to geranic acid or citronellic acid (Fig. 6d-f) and the total amount of exogenous geraniol decreased in crude lysate only when NAD^+^ was present (Fig. S10), we suspected that this NAD-dependent aldehyde dehydrogenase may be responsible. Thus, we targeted CgI_2844 to generate strains Δ*2844* (BL-8) and Δ*adhC* Δ*2844* (BL-9) to determine if this enzyme was responsible for the oxidation of citral or citronellal or misannotated and instead had function as an aldehyde reductase. However, loss of CgI_2844 did not impact the biotransformation of geraniol and citral to geranic acid or citronellic acid in Δ*adhC* Δ*2844* (Fig. S12). While AdhC reduces citral and citronellal to their respective alcohols, the aldehyde dehydrogenase(s) oxidizing citral and citronellal, alcohol dehydrogenase(s), and secondary aldehyde reductase remain unknown (Fig. 6g).

## 4. Conclusion

Monoterpenes have tremendous utility to various industries including as flavors and fragrances, though titers in conventional GRAS industrial microbes have been exceedingly low thus far. This study demonstrates the highest monoterpene production achieved in *C. glutamicum* to date at more than 700 mg/L, an important step in developing this organism as an industrial GRAS monoterpene platform. We leveraged an episomal mevalonate pathway and genetic modifications to increase precursor flux while reducing volatility of the monoterpene products through the use of a biphasic production system. Future engineering efforts can now leverage this mevalonate-based strategy to produce the entire suite of isoprenoid natural products in *C. glutamicum,* though additional enzyme and strain optimization will likely be required.

We also demonstrated that a putative aldehyde reductase, AdhC, possesses additional substrate promiscuity for acyclic aldehydes. However, we were unable to identify any other enzymes in *C. glutamicum* involved in the oxidation or reduction of geraniol, an avenue that remains open for further investigation.

## Author Statement

**Conceptualization**: B.A.L; **Data Curation:** B.A.L, A.N.P, and Y.C.; **Formal Analysis:** B.A.L, A.N.P, Y.C., Y.L.; **Methodology:** B.A.L, Y.C., Y.L., L.E.V.; **Investigation**: B.A.L, M.K., A.N.P, Y.C., Y.L., L.E.V, A.C.R., G.A.H, X.B.T., and B.W.; **Software:** A.N.P and Y.C.; **Visualization:** B.A.L, A.N.P., and Y.L.; **Validation:** B.A.L, A.N.P, and Y.C.; **Writing - original draft:** B.A.L, Y.C., and A.N.P.; **Funding Acquisition, Resources, and Supervision:** C.J.P and J.D.K; **Writing - review & editing:** All authors

## Supporting information

SuppFigures

## Acknowledgements

We would like to sincerely thank Prof. Dr. Volker Wendisch for providing plasmids, Dr. Thomas Eng and Dr. Aparajitha Srinivasan for their advice regarding the production of isoprenol, and Dr. David Carruthers and Dr. Jinho Kim for their assistance with GC-MS/GC-FID analysis. This work was funded by the DOE Joint BioEnergy Institute (https://www.jbei.org) supported by the U.S. Department of Energy, Office of Science, Office of Biological and Environmental Research through contract DE-AC02-05CH11231 between Lawrence Berkeley National Laboratory, the U.S. Department of Energy, a Department of Energy Office of Science Distinguished Scientist Award to J.D.K., and the Philomathia Foundation.

## Data Availability

Data access is included in the manuscript.

## Appendix A. Supplementary Data

The following is the supplementary data for this article:

