## Supplementary material for "Development of *Corynebacterium glutamicum* as a monoterpene production platform": SuppFigures

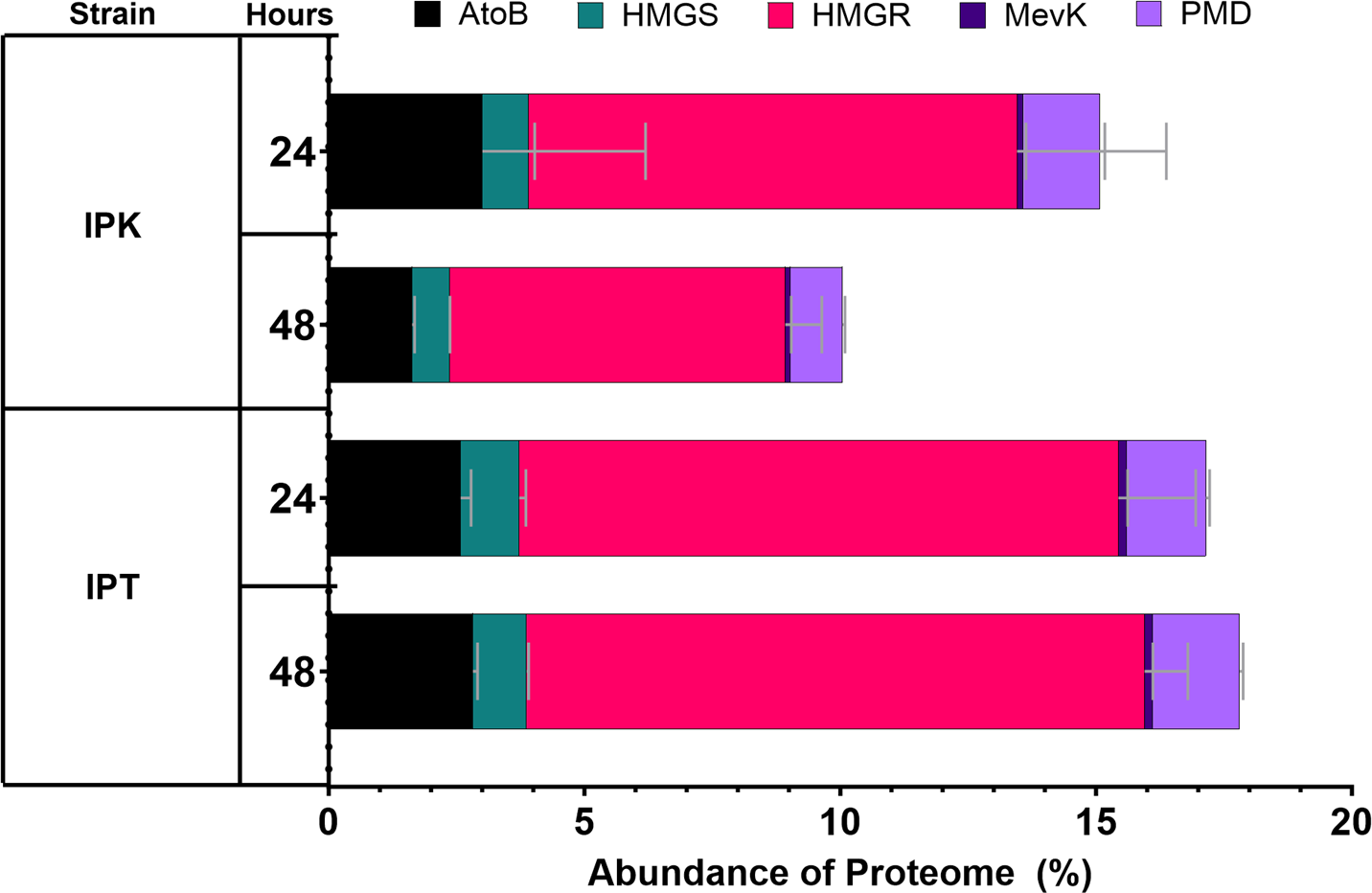
**A**

**B**

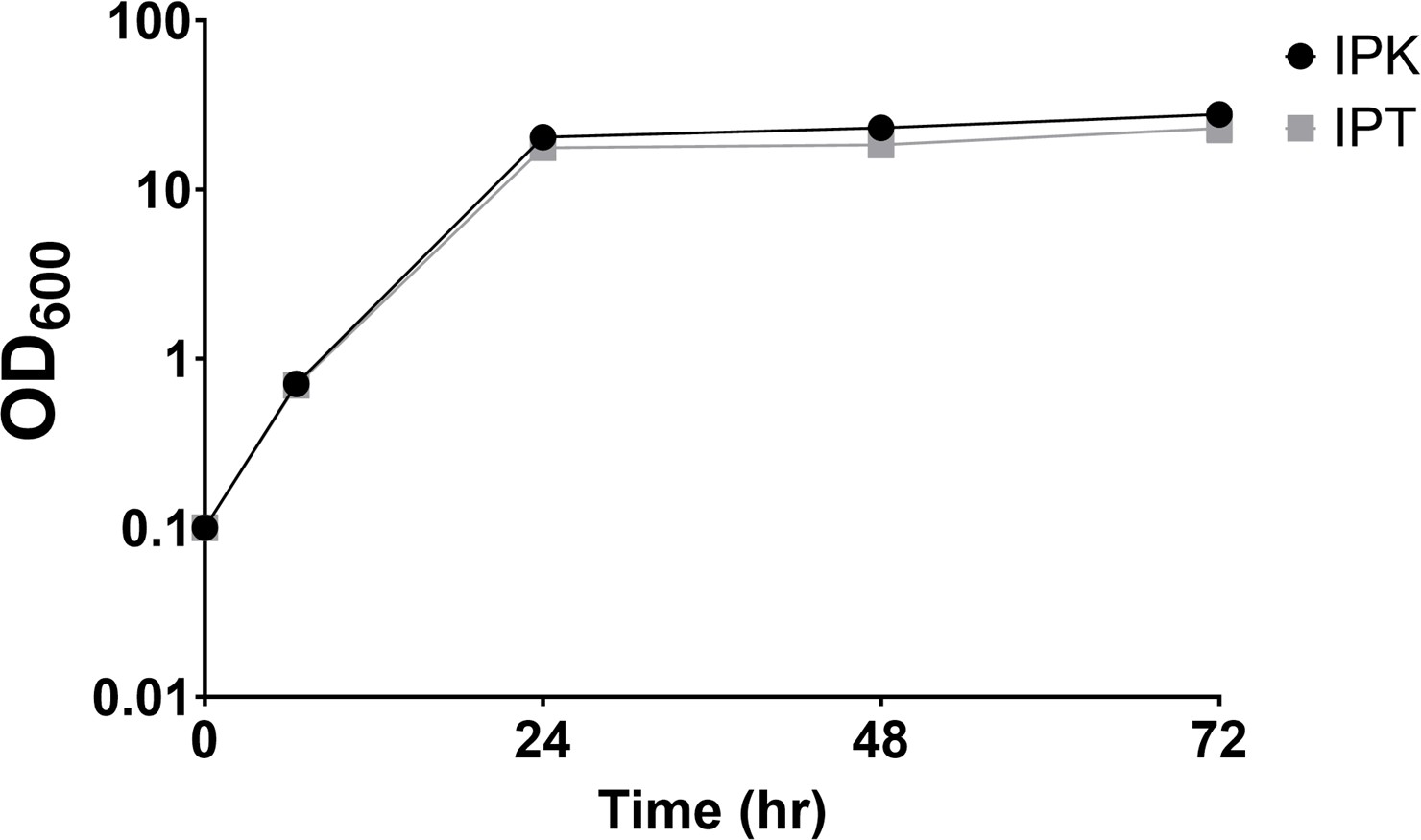

**Figure S1. Pathway protein expression and growth of isoprenol production strains.** The IPK and IPT strains were induced at an OD_600_ of 0.8 and harvested at the indicated times following induction to measure pathway protein expression (a) and growth (b).

**Table S1**. Further information for *C. glutamicum* JBEI 1.1.2 genes targeted in this study

| **Gene ID** | **Locus Tag** | **Genbank Accession** | **Gene Product Name** | **ATCC 13032**  **homolog (Protein Identity)** | **Genome Coordinates** |
| --- | --- | --- | --- | --- | --- |
| 2821588  608 | Ga0373873_ 1733 | NII87708.1 | NADPH-dependent 2,4-dienoyl-CoA reductase/sulfur reductase-like enzyme | NCgl2615 (29%) | 1869784..187089  9 (+) |
| 2821589  183 | Ga0373873_ 2308 | NII88272.1 | NADPH-dependent 2,4-dienoyl-CoA reductase/sulfur reductase-like enzyme | creE, NCgl0525 (98%) | 2466245..246751  3 (+) |
| 2821589  274 | Ga0373873_ 2399 | NII88357.1 | geranylgeranyl reductase family protein | NCgl0455 (99%) | 2551886..255315  1 (+) |
| 2821589  729 | Ga0373873_ 2854 | NII88801.1 | NADPH-dependent 2,4-dienoyl-CoA reductase/sulfur reductase-like enzyme | NCgl2615 (37%) | 3003695..300487  3 (-) |
| 2821589  963 | Ga0373873_ 3088 | NII89023.1 | 2,4-dienoyl-CoA reductase-like NADH- dependent reductase (Old Yellow Enzyme family) | NCgl2942 (98%) | 3243429..324455  0 (-) |
| 2821589  967 | Ga0373873_ 3092 | NII89027.1 | 2,4-dienoyl-CoA reductase-like NADH- dependent reductase (Old Yellow Enzyme family) | NCgl2938 (99%) | 3247242..324835  4 (+) |
| 2821587  100 | Ga0373873_ 225 | NII86254.1 | NADPH-dependent 2,4-dienoyl-CoA reductase/sulfur reductase-like enzyme | NCgl2615 (98%) | 268090..269325 (+) |
| 2821589  459 | Ga0373873_ 2584 | NII88541.1 | uncharacterized zinc-type alcohol dehydrogenase-like protein | adhC/FudC, NCgl0324 (98%) | 2734177..273523  8 (+) |
| 2821587  014 | Ga0373873_ 139 | NII86168.1 | propanol-preferring alcohol dehydrogenase (adhA) | adhA, NCgl2709 (99%) | 164893..165930 (+) |
| 2821587  198 | Ga0373873_ 323 | NII86351.1 | pyruvate dehydrogenase (quinone) | poxB, NCgl2521 (99%) | 374338..376077 (+) |
| 2821586  897 | Ga0373873_ 22 | NII86052.1 | L-lactate dehydrogenase | ldhA, NCgl2810 (99%) | 22587..23531 (+) |
| 2821589  067 | Ga0373873_ 2192 | NII88156.1 | geranylgeranyl diphosphate synthase type II | crtE, NCgl0600 (95%) | 2340971..234207  7 (+) |
| 2821587  696 | Ga0373873_ 821 | NII86822.1 | geranylgeranyl diphosphate synthase type I | idsA, NCgl2092 (99%) | 897151..898251 (-) |
| 2821589  719 | Ga0373873_ 2844 | NII88791.  1 | acyl-CoA reductase-like NAD-dependent aldehyde dehydrogenase | betB, NCgl0523 (39%) | 2993772..2995271 (-) |

**Table S2.** Screening primers

| **Primer Name** | **Sequence (5’ → 3’)** |
| --- | --- |
| crtEscreen_F | agtagcagccattccccag |
| crtEscreen_R | cacgggctctttcaggttc |
| idsAscreen_F | acggcaaggtcacagaattg |
| idsAscreen_R | ccgaggtcgacgatgagg |
| 225screen_F | tttgggatcacgttccctt |
| 225screen_R | gaactgcagccaaaccatta |
| 1733screen_F | ccaagaattgggaatggga |
| 1733screen_R | tgtgcagatccatagaaactatt |
| 2308screen_F | atgcctagtccacgcactgt |
| 2308screen_R | tgtcttcgccgtctaacca |
| 2399screen_F | gttaagtggtggattacgggg |
| 2399screen_R | gtatggattgtctttggaaggc |
| 2854screen_F | gttgctgtaatcgctcccgt |
| 2854screen_R | aaggagattgctggattgcc |
| 3088screen_F | gcggttggaggtaaagcct |
| 3088screen_R | cgaaacaaagaggccatcag |
| 3092screen_F | cttggttccgcagaactgaata |
| 3092screen_R | tgactcgctcctgccgtc |
| adhCscreen_F | cgaatcctacgatgaaaacg |
| adhCscreen_R | aacgctgacatcggaatcat |
| adhAscreen_F | ccatcatgagccgttgatta |
| adhAscreen_R | gtgctcgaatgacatctccg |
| 2844screen_F | ctcgaacgtgaacaagacca |
| 2844screen_R | aatcagatcgttctgcccac |

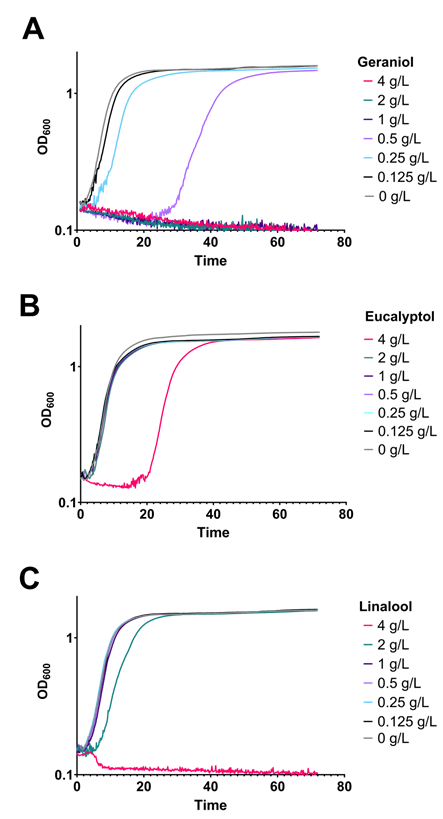

**Figure S2. Monoterpene toxicity growth curves of *C. glutamicum*.** *C. glutamicum* was incubated in CGXII with the addition of (a) geraniol, (b) eucalyptol or (c) linalool.

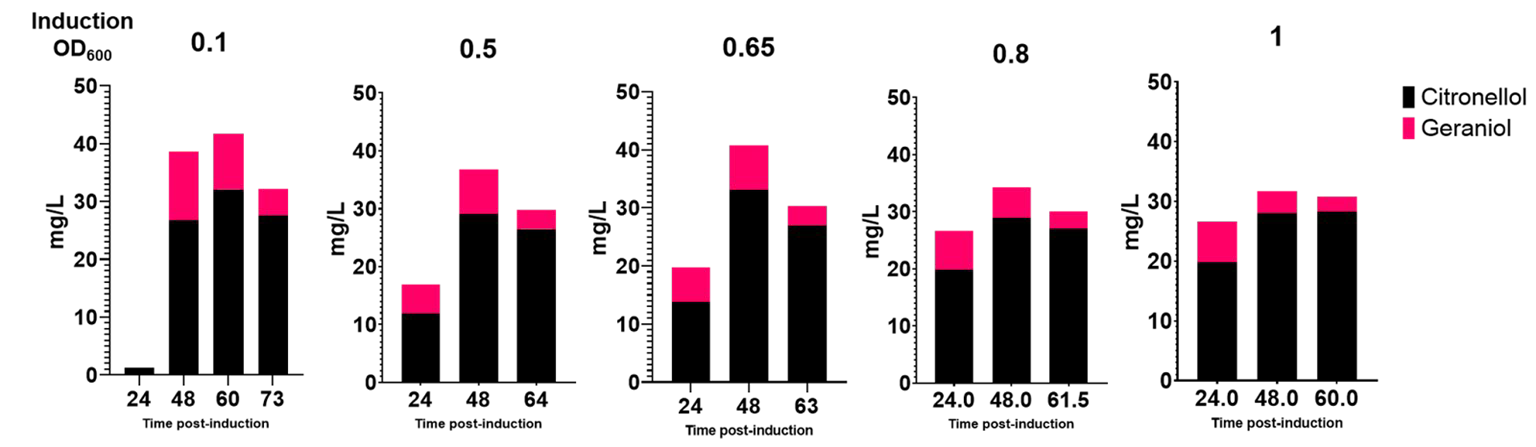

**Fig. S3. Induction optimization of monoterpene production.** *C. glutamicum* cultures of PL-114 were induced at the indicated optical density.

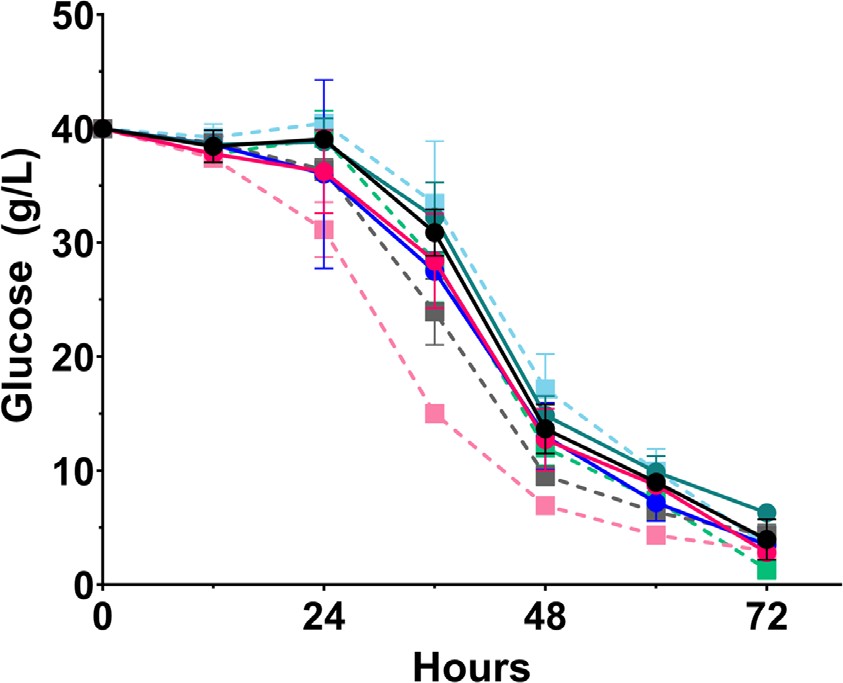
**A**

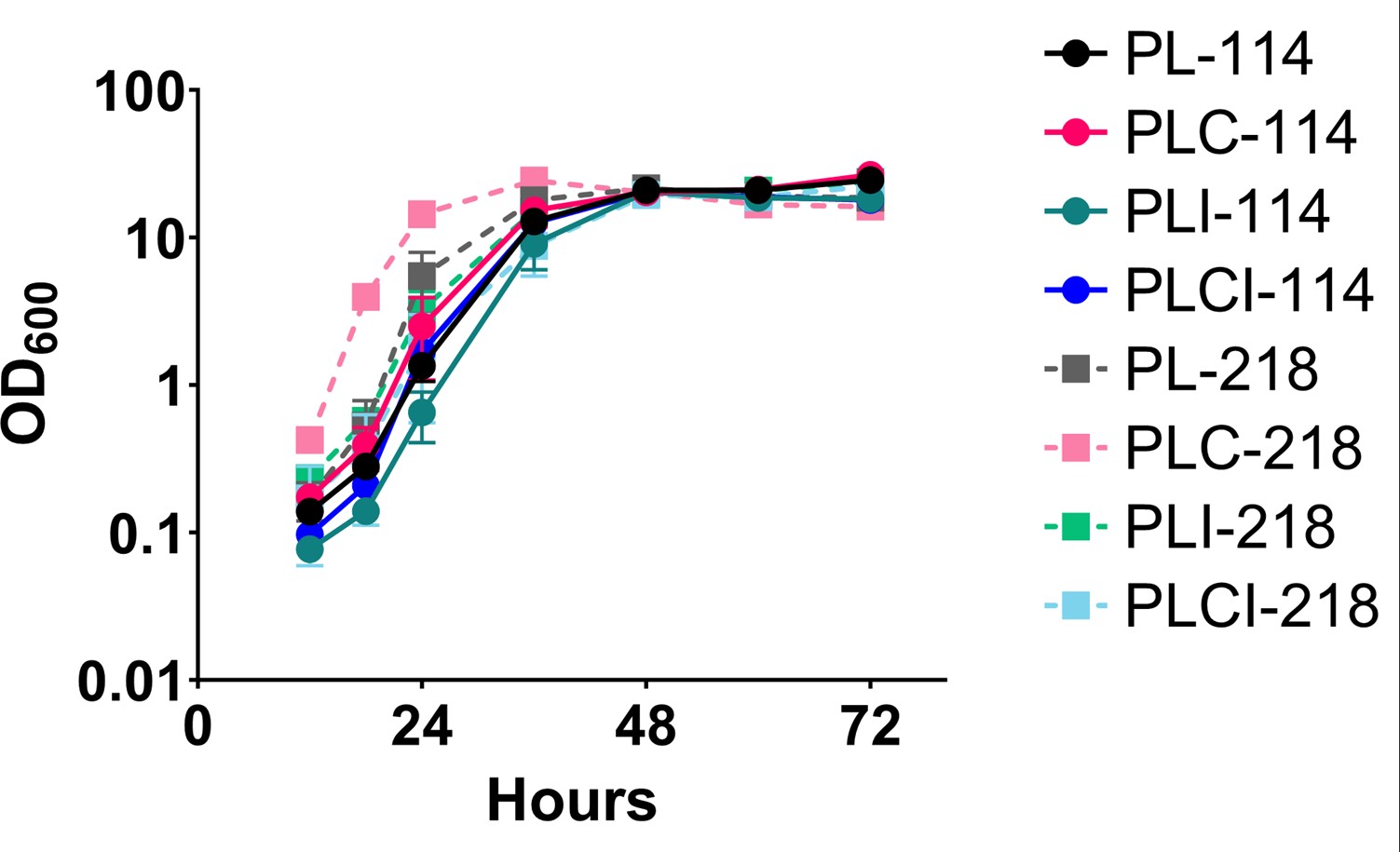

**B**

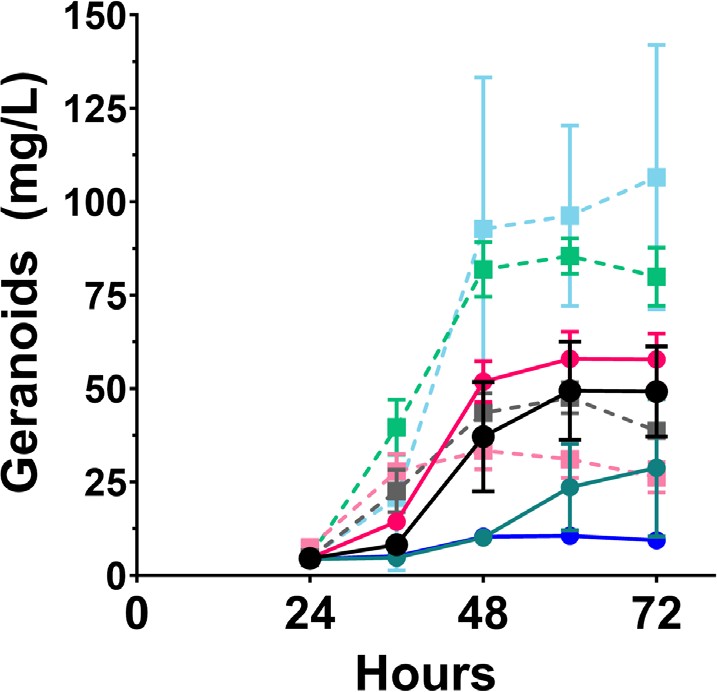
**C**

**Figure S4. Kinetic metrics of geranoid production strains.** Glucose consumption (a), optical density (b), and geranoid production over 72 hours was measured for each strain grown in 5 mL CGXII containing 40 g/L glucose inoculated and induced with 0.5 mM IPTG at an OD_600_ of 0.1.

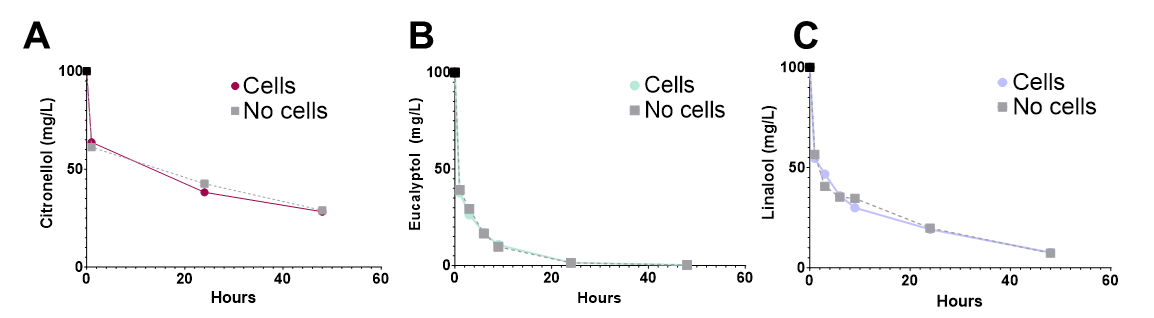

**Figure S5. Monoterpene volatility and catabolism.** Assessing citronellol (a), eucalyptol (b), and linalool

(c) volatility and catabolism by *C. glutamicum.* The black data point indicates the initial calculated concentration added to the medium and was not measured by GC-MS.

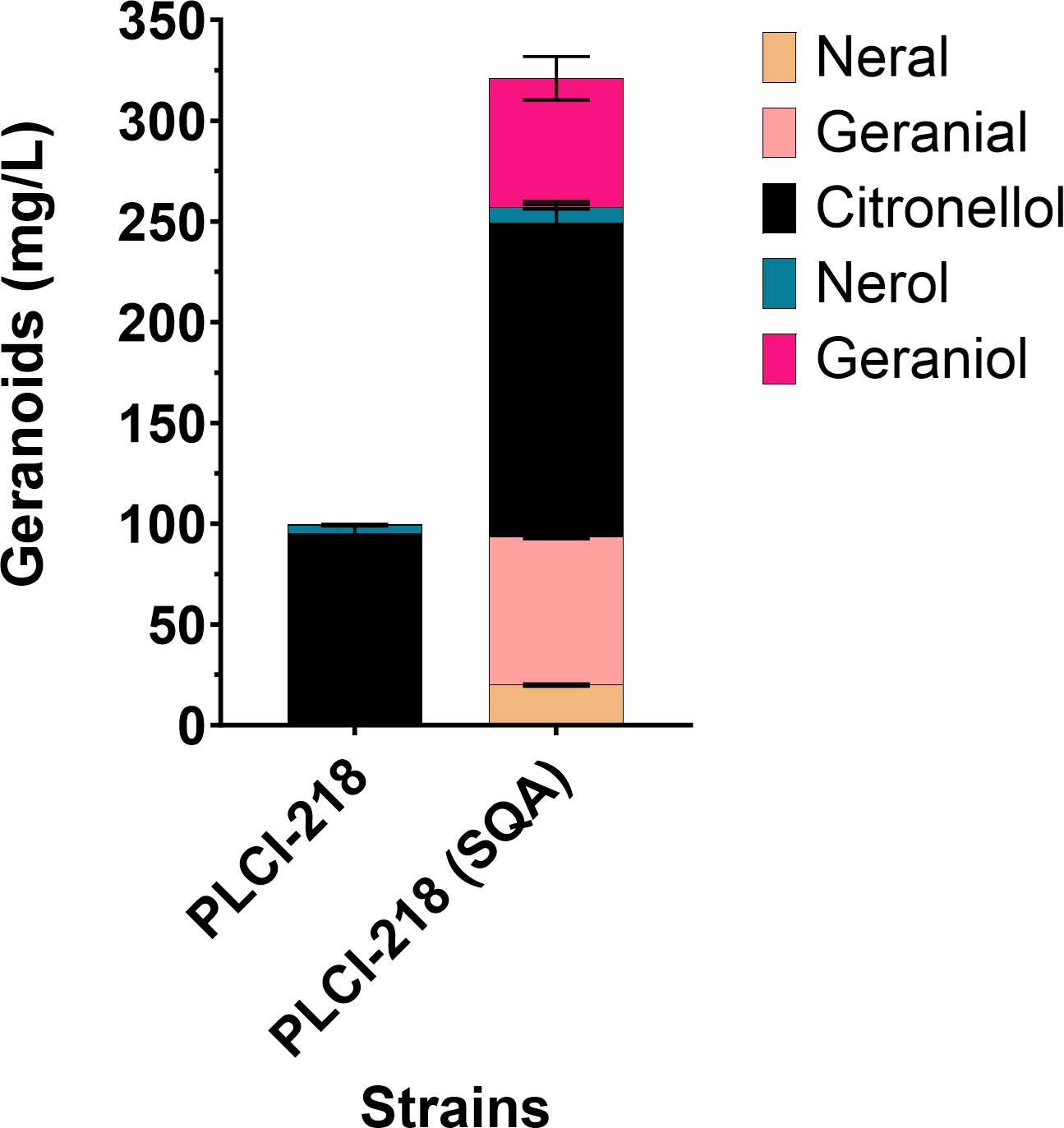

**Figure S6. Geranoid composition produced by PLCI-218 using a 10% squalane overlay.** Composition of geranoids measured by GC-MS after 72 hours with use of a 10% squalane overlay added 12 hours following induction.

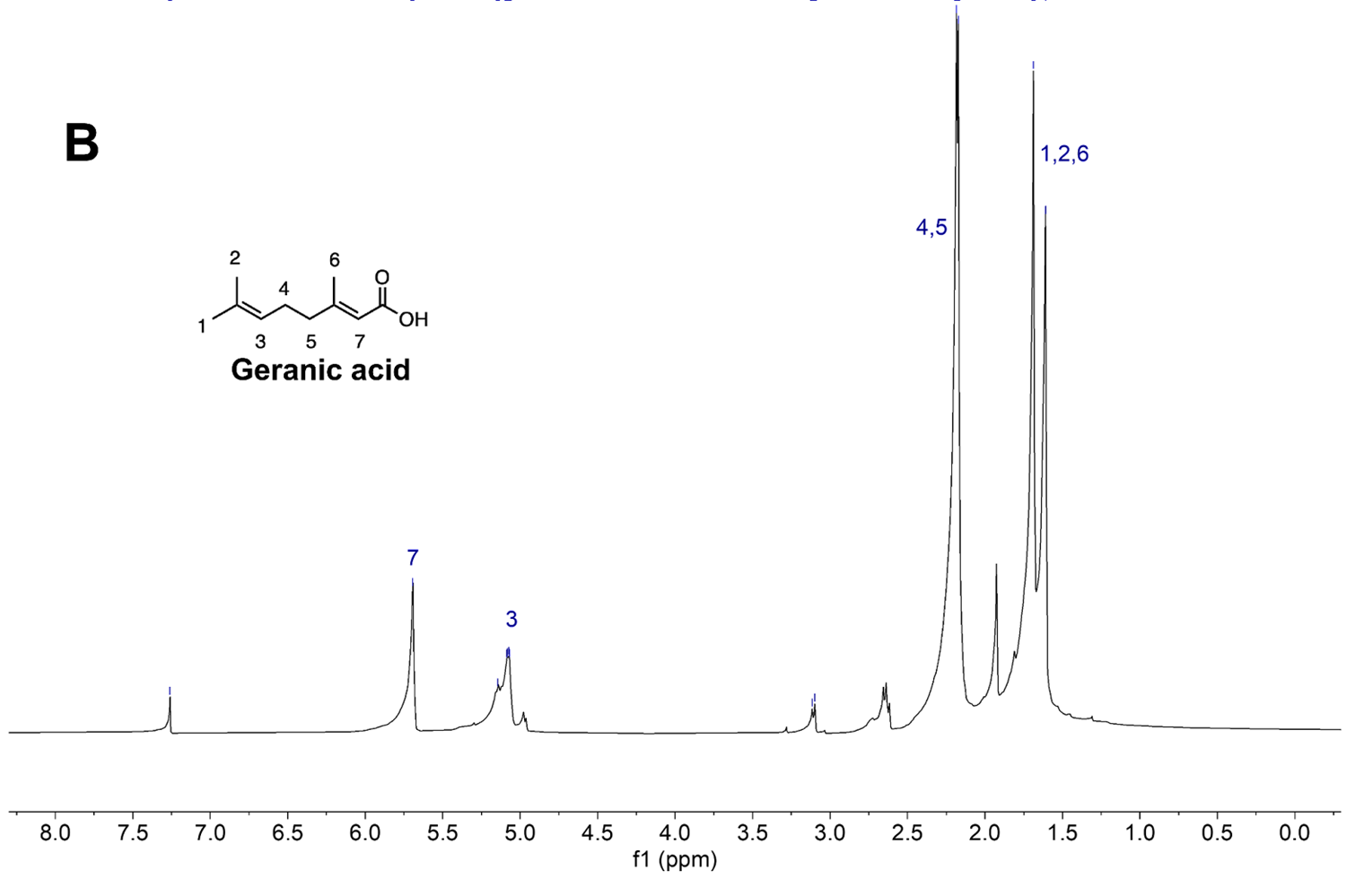

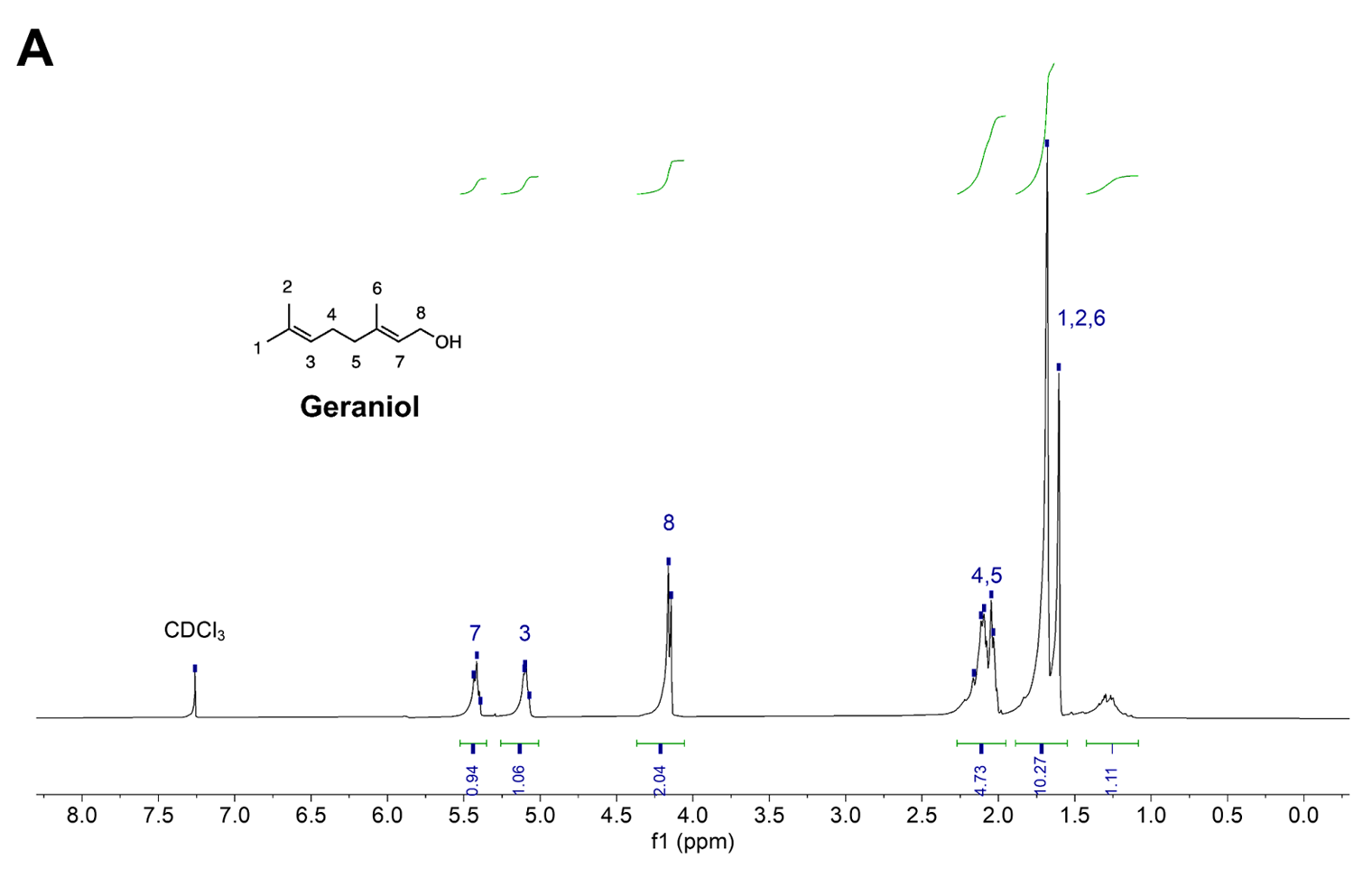

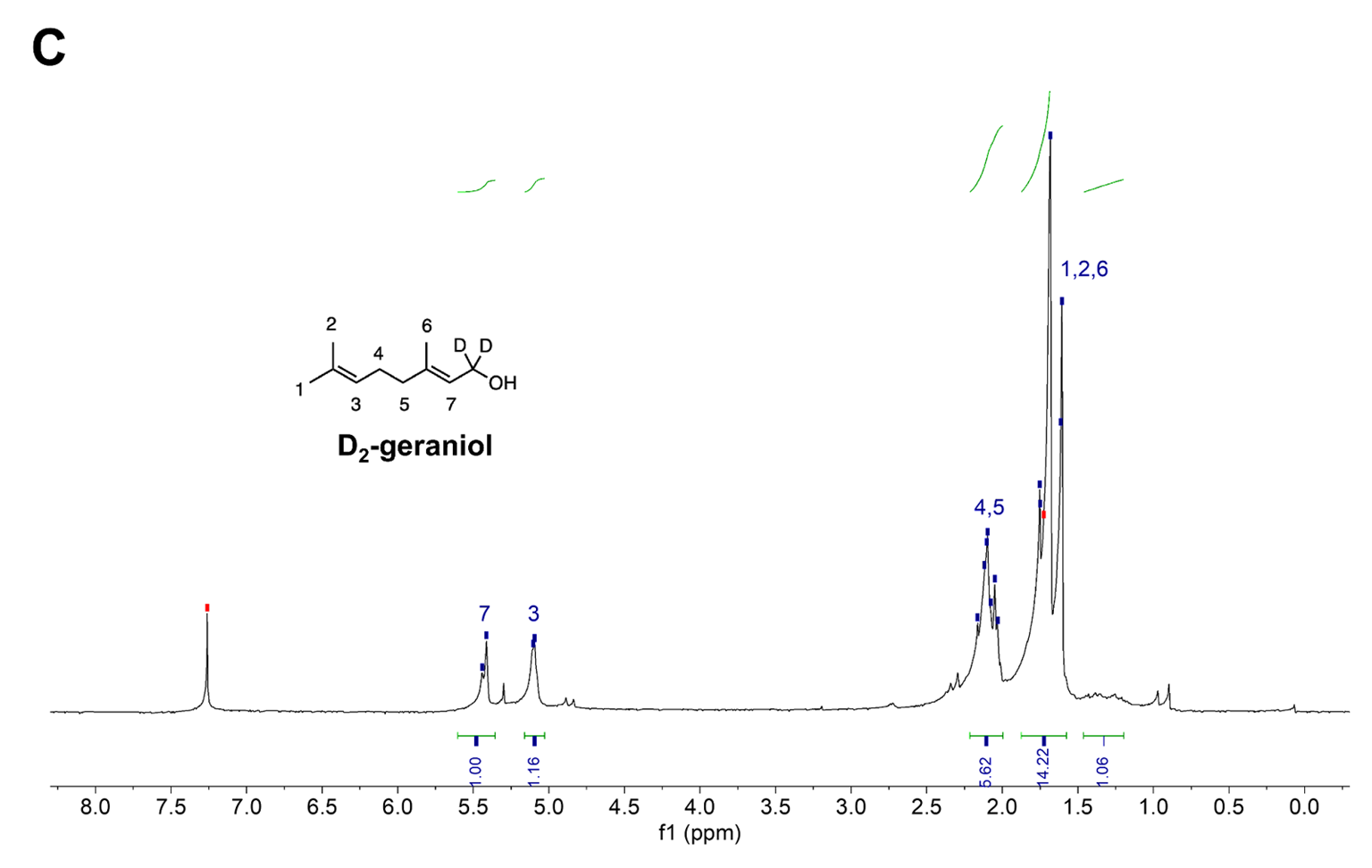

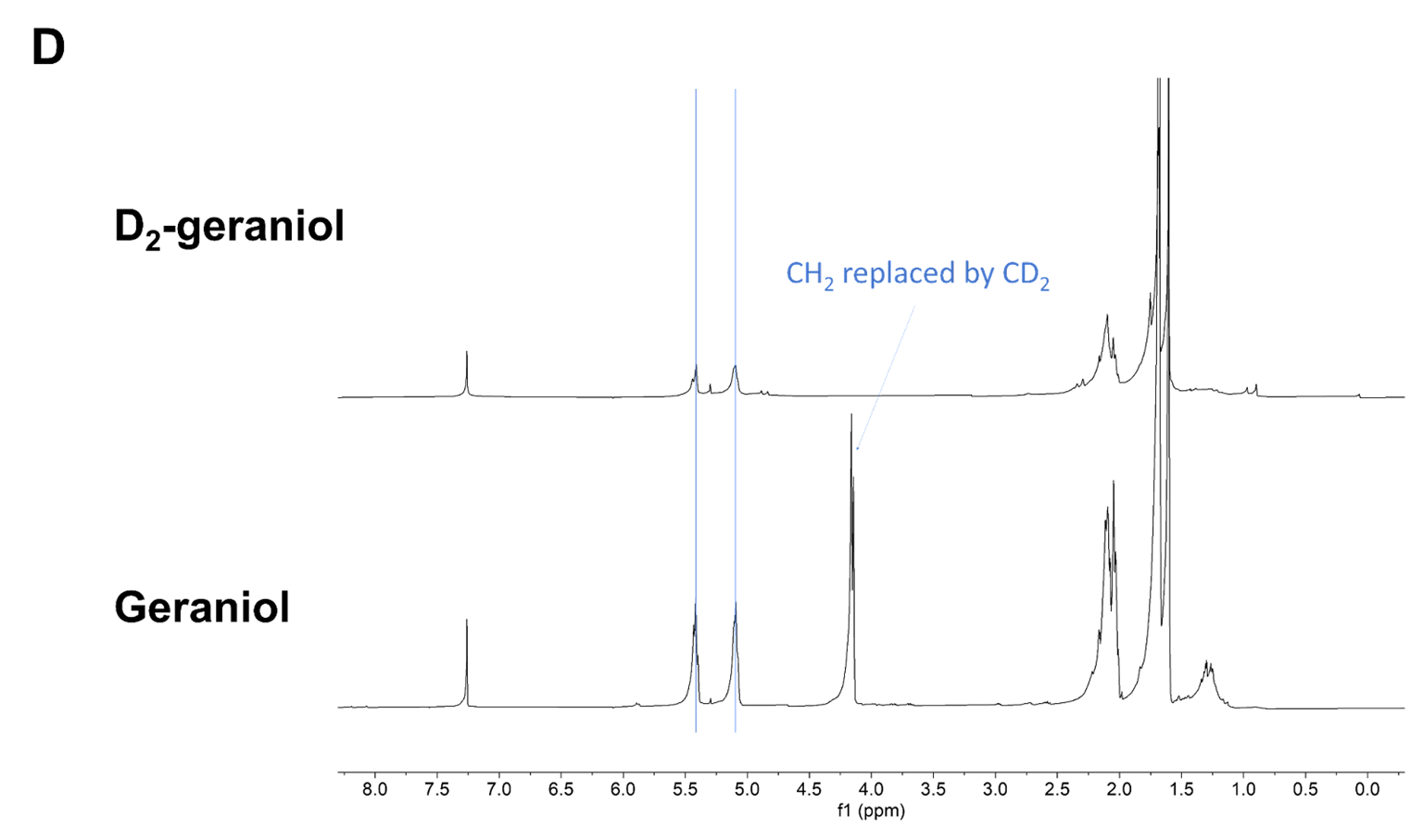

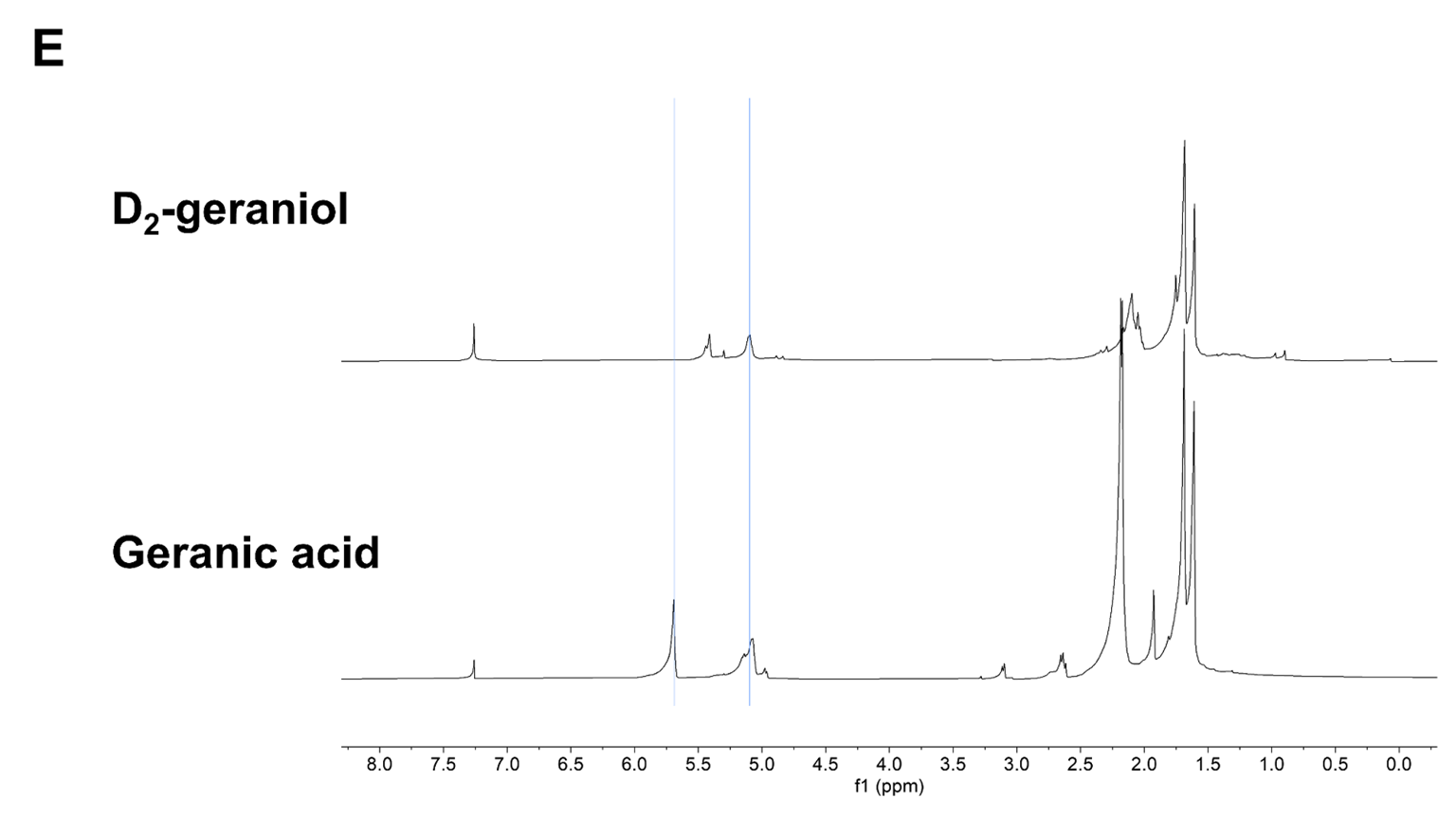

**Figure S7. ^1^H NMR of synthesized D_2_-geraniol from geranic acid and LiAlD_4_.** ^1^H NMR of geraniol (a), geranic acid (b), and D_2_-geraniol (c). The spectra are compared in e to confirm deuteration of geraniol (d) and complete conversion of geranic acid (e).

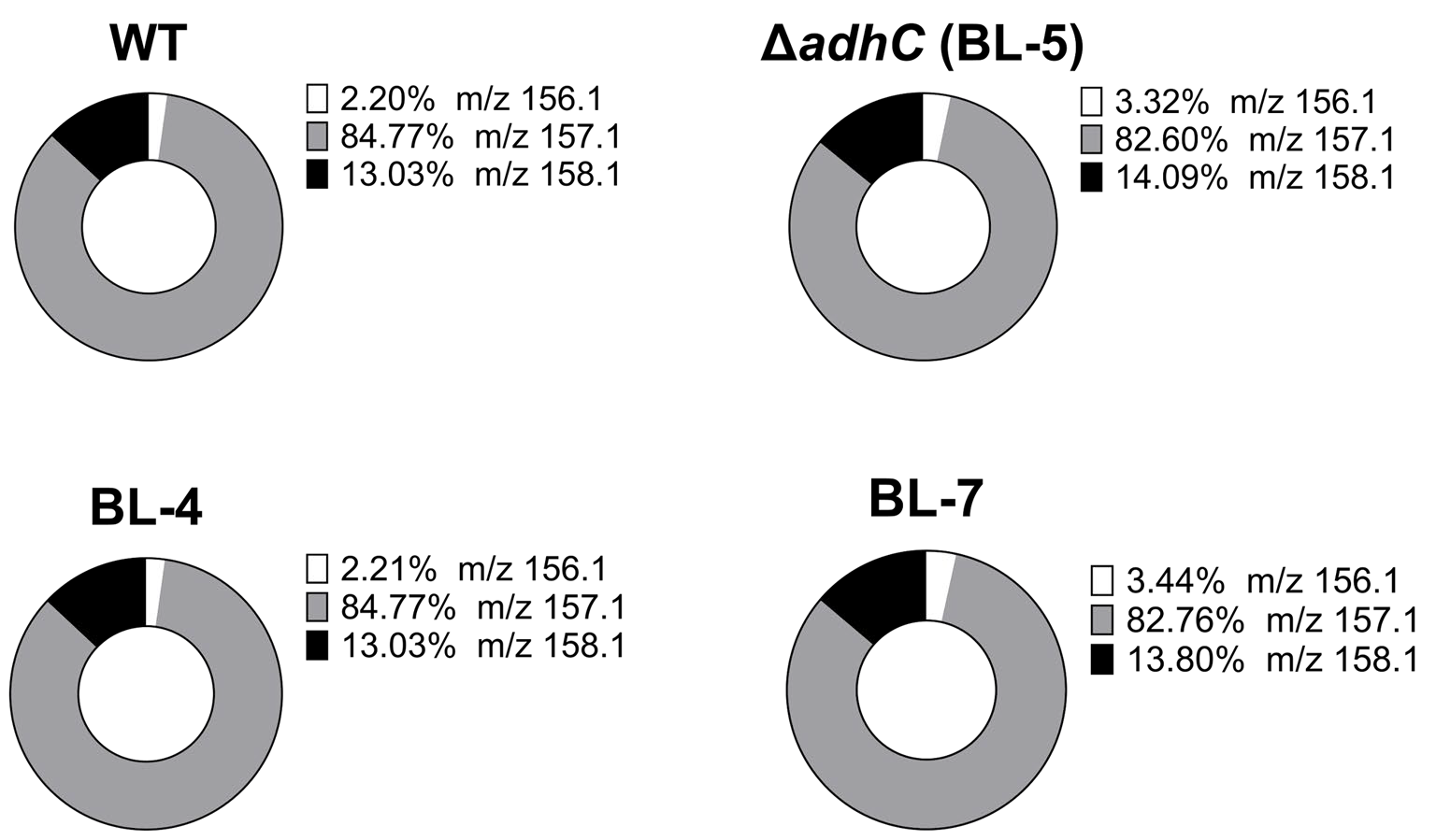

**Figure S8. Ratios of deuterium-labeled and unlabeled citronellol following biotransformation of D_2_- geraniol by *C. glutamicum*.** The indicated strains were incubated in CGXII in the presence of D_2_-geraniol for 24 hours and evaluated by GC-MS.

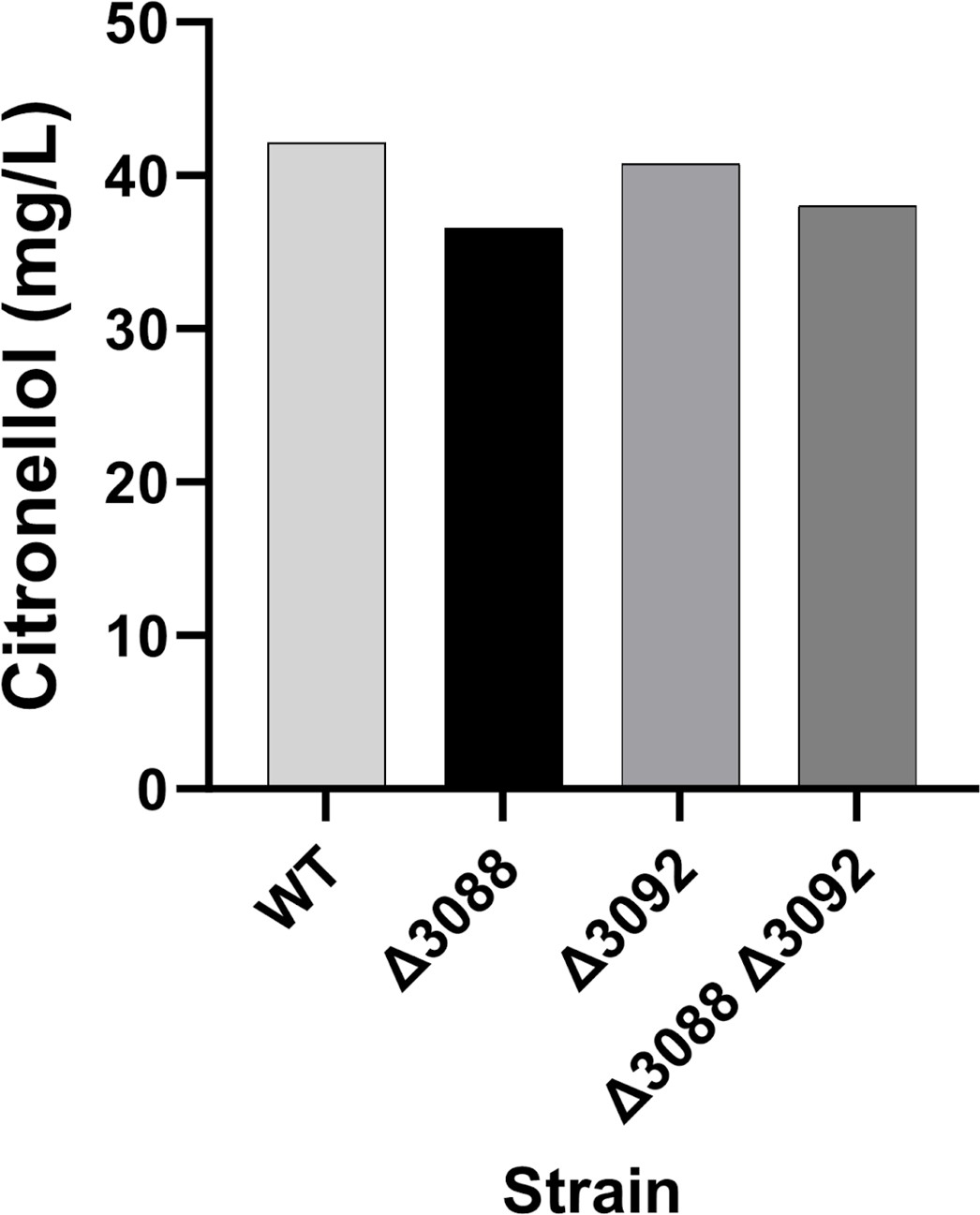

**Figure S9. Biotransformation of geraniol to citronellol following the disruption of two old yellow enzyme reductases.** The indicated strains were incubated in the presence of geraniol for 24 hours to determine their capacity for reducing geraniol to citronellol.

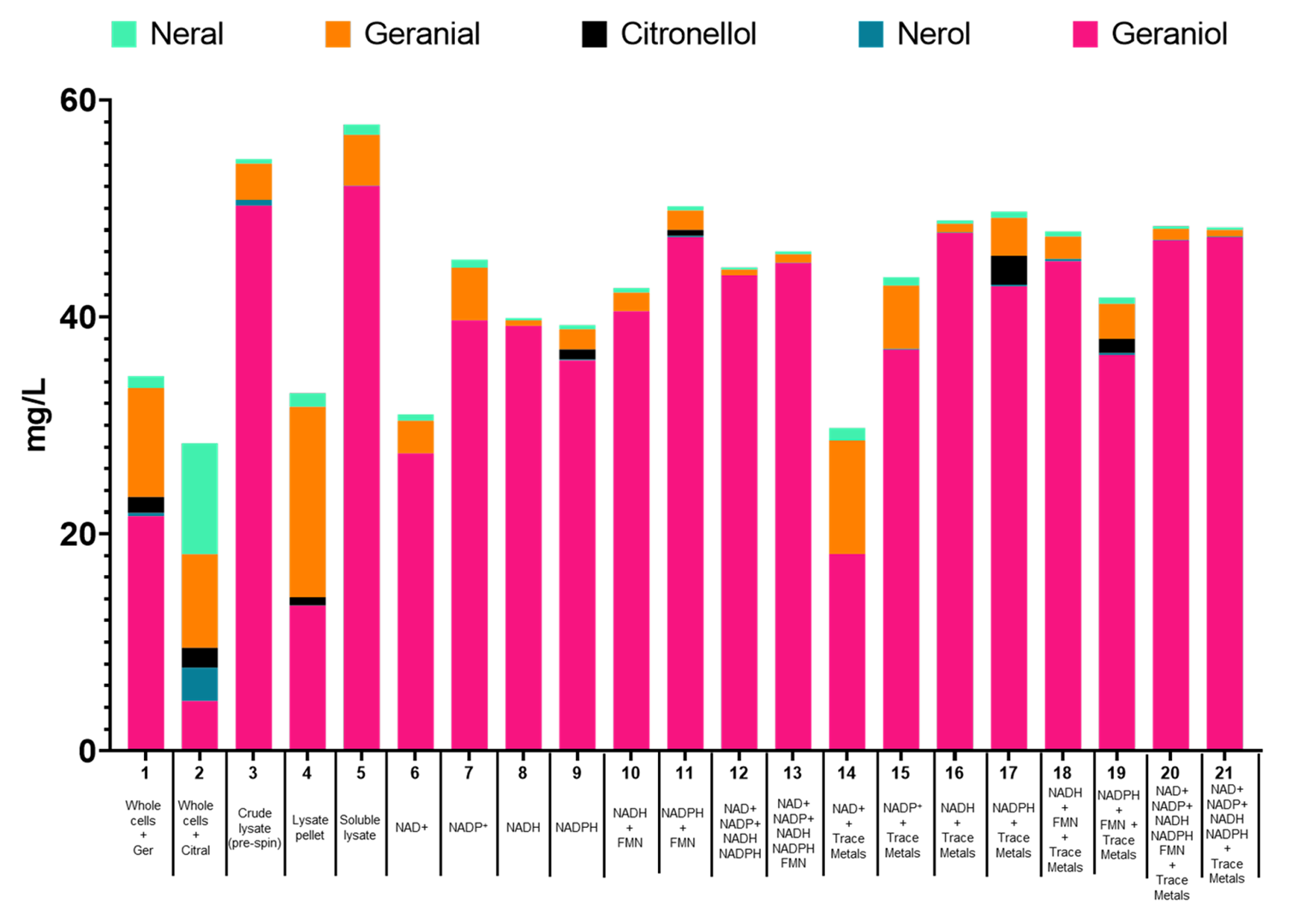
**Figure S10. Crude lysate assay to determine cofactor requirements for geraniol reduction.** The soluble fraction of a crude cell lysate (and aliquots at different steps during its preparation) were incubated with the indicated cofactors and trace metals to determine cofactor requirements for conversion of geraniol/citral to citronellol. Lane 2 was incubated with citral and all other lanes were incubated with geraniol.

**A**

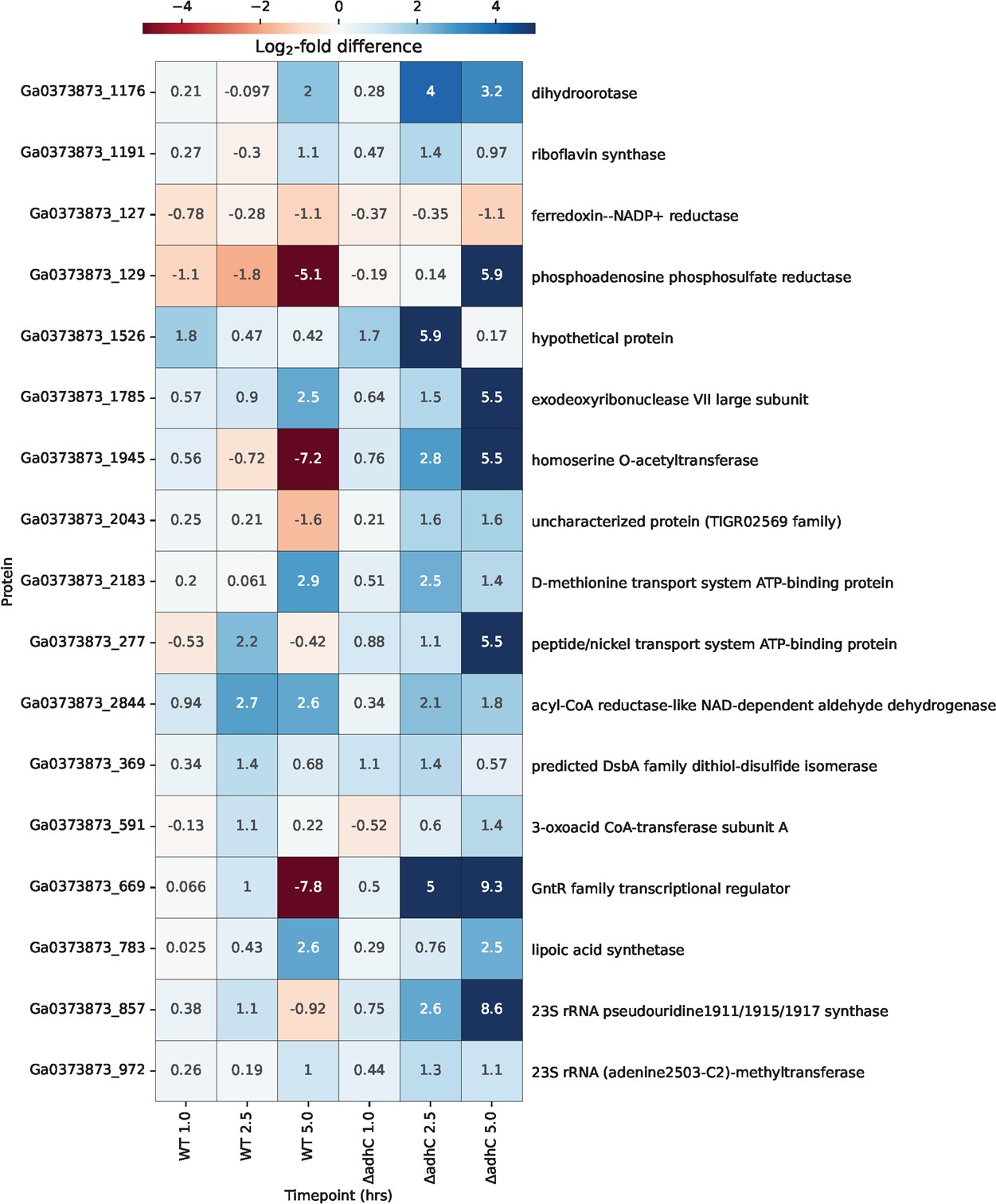

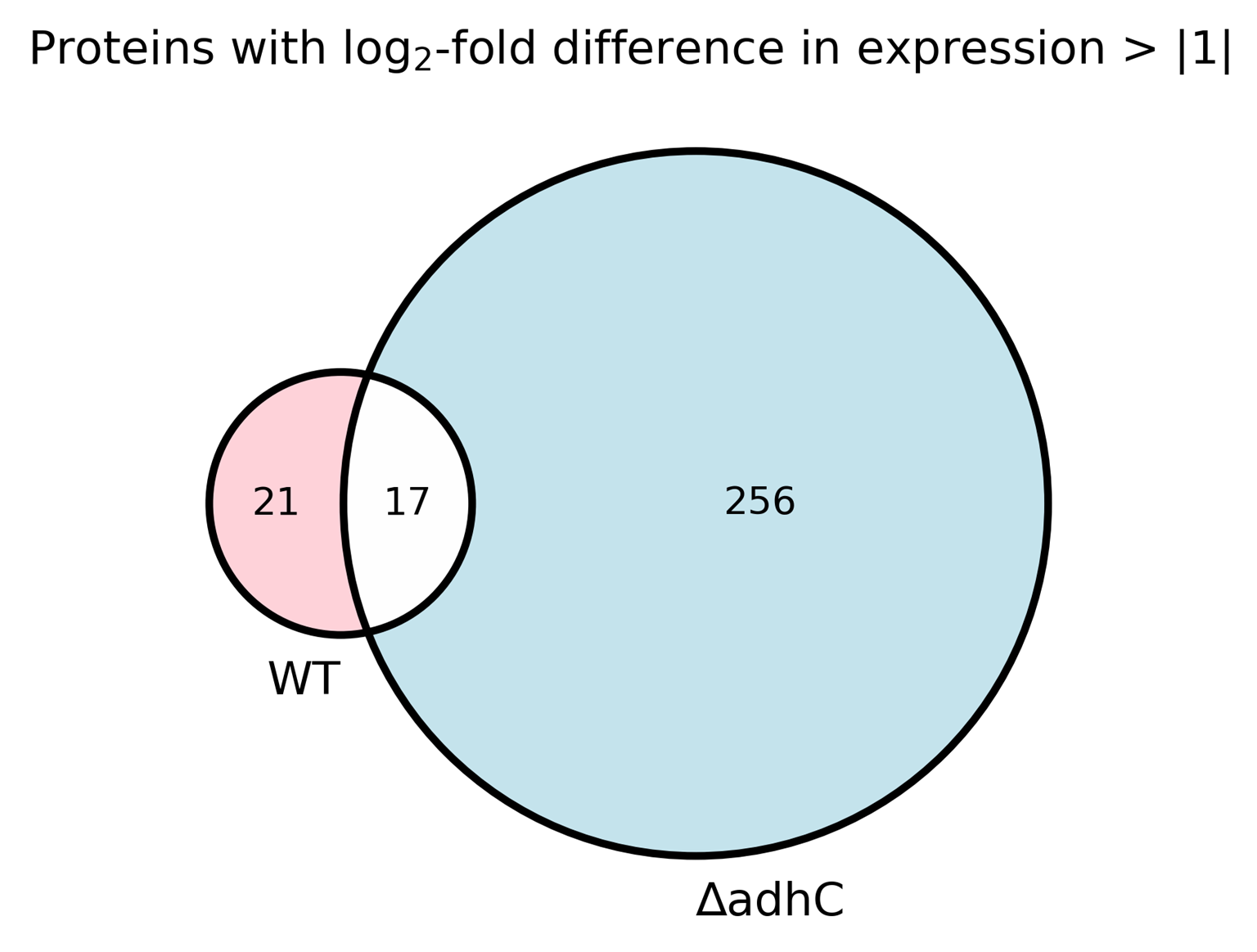

**B**

**Fig. S11. Citral-responsive proteins differentially expressed in WT and ∆*adhC.*** (a) The total number of unique and shared proteins in each strain that exhibited at least a log_2_-fold difference in expression > |1| in at least one time point after exposure to 100 mg/L citral vs. ethanol. (b) The 17 citral-responsive proteins shared between the strains are listed with the corresponding log_2_-fold difference for each time point

measured.

**B**

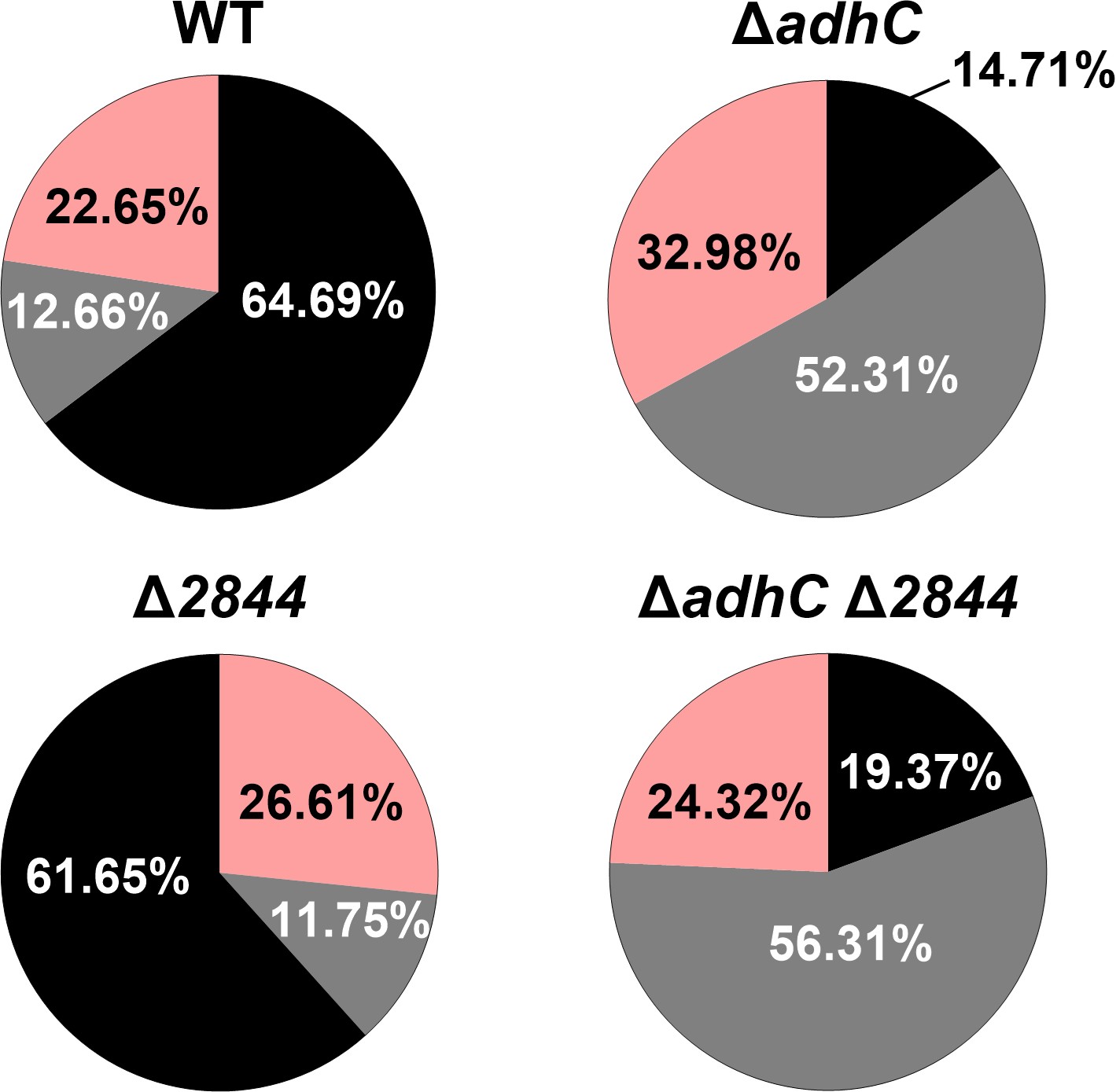

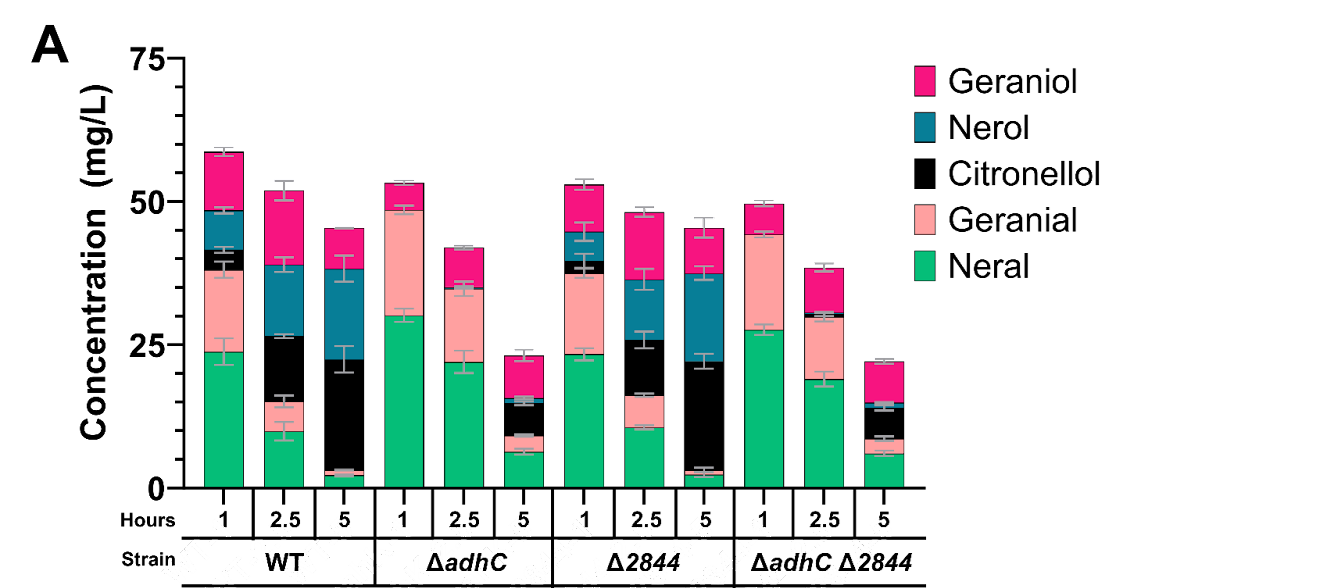

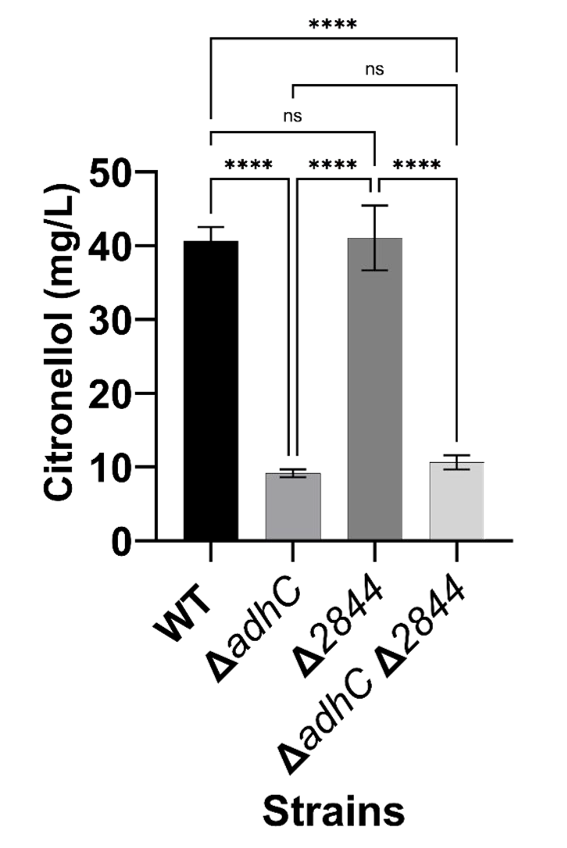

**C	D**

**E	F**

**Figure S12.** Evaluating the cooperative disruption of a putative aldehyde reductase and aldehyde dehydrogenase on the conversion of citral and geraniol. Citral (a) or geraniol (b-f) were incubated with the indicated strains in BHI (a) or CGXII (b-f) and the biotransformed products were analyzed by GC-MS at the indicated time (a) or after 24 hours (b-f). For panel c, an ordinary one-way ANOVA with multiple comparisons was performed where p < 0.0001.
